# Topoisomerase IIβ Kinase: A protein processed from HIV-1 envelope by cysteine proteases

**DOI:** 10.64898/2026.10.26.761857

**Authors:** Hema Verma, Satyajit Mukhopadhyay, Lokeswara S Balakrishna, Ritika Das, Abhash Kumar, Sangeetha Balasubramanian, Anand K Kondapi

**Author notes:** Department of Biochemistry and Molecular Biology, Central University of Kerala, Tejaswini Hills, Periye (PO), Kasaragod (DT), Kerala-671316, India. Equally contributed (Hema Verma, Satyajit Mukhopadhyay, Lokeswara S Balakrishna). Anand K Kondapi **Email:**.

## Abstract

Topoisomerase IIβ (TopoIIβ) is activated by TopoIIβ kinase (TopoIIβK) during HIV-1 infection. This study demonstrates that TopoIIβK is produced within 15-30 minutes post-infection and is conserved across the major HIV-1 subtypes (A, B, C, and D), although its formation kinetics vary among subtypes. Immunological mapping using specific anti-gp120 antibodies suggests that TopoIIβK is derived from the N-terminal region of gp120. Notably, TopoIIβK co-localizes with endolysosomes shortly after viral entry. Inhibition of cysteine proteases using E64d and Z-FY-CHO abrogates TopoIIβK production in both HIV-1-infected and envelope-transfected cells. These findings suggest that TopoIIβK is a conserved polypeptide product of gp120 processing within endolysosomes during the early stages of HIV-1 infection, highlighting a potentially conserved mechanism that regulates early viral infection across major HIV-1 subtypes.

## Introduction

HIV-1, the etiological agent of acquired immunodeficiency syndrome (AIDS), propagates by hijacking the cellular machinery of host CD4+ T lymphocytes [1]. HIV-1 entry into host cells is mediated by the binding of gp120 to CD4 receptors, followed by co-receptor engagement and subsequent membrane fusion mediated by the gp41 domain through the formation of a six-helix bundle [2]. Several studies have also reported the involvement of the endosomal pathway in viral entry [3]. In a study employing siRNA-mediated knockdown of Rab5, Rab7, and Rab11, the endosomal and proteasomal pathways were implicated in viral entry and replication, suggesting that multiple cellular pathways may contribute to these processes [4].

Among the cellular factors usurped to support the viral life cycle, host DNA Topoisomerases play a significant role in viral replication [5,6]. Among the isoforms, Topoisomerase IIβ (TopoIIβ) is involved in reverse transcription [7,8,9] and Tat-mediated transcription [10]. TopoII isoforms exhibit differential dynamics of phosphorylation, with TopoIIβ phosphorylating activity (TopoIIβK) appearing at 1-2 hours post-infection, followed by TopoIIα phosphorylating activity at 4 hours post-infection [11]. TopoIIβK activity was purified from HIV-1 particles [11]. Further, TopoIIα kinase was sensitive to PD9859, a MAP kinase inhibitor, while TopoIIβ kinase was resistant to known kinase inhibitors [11,12]. Screening studies using several pyridines and coumarin derivatives yielded potent inhibitors targeting TopoIIβK that exhibited significant inhibition of HIV-1 infection [13,14]. The objective of this communication is to unravel the origin of TopoIIβK.

## Materials and Methods

### Materials

The cell lines, viral isolates, gp120 envelope clones, and anti-gp120 antibodies used in this study were obtained from the NIH HIV Reagent Program. Detailed information is provided in Supplementary Tables S1–S4.

Peripheral blood mononuclear cells (PBMCs) were isolated from healthy donors in accordance with the approved Institutional Ethics Committee (IEC) protocol and cryopreserved in liquid nitrogen. Following revival, PBMCs exhibiting 60%–70% viability were used for virus propagation and kinetic studies.

HIV-1 NL4-3 (subtype B; Cat. No. ARP-3420) and HIV-1 isolates representing subtypes A (99KE_KNH1088), B (90US_873), C (94IN_20635-4), D (98UG_57128), and circulating recombinant form CRF_AE (96TH_M02138) (Supplementary Table S1), all belonging to group M, were obtained from the NIH HIV Reagent Program, Division of AIDS, National Institute of Allergy and Infectious Diseases (NIAID), National Institutes of Health (NIH), USA. HIV-1 gp120 envelope clones (Supplementary Table S2) and anti-gp120 monoclonal antibodies (Supplementary Table S3) were also obtained from the same source.

A mouse monoclonal antibody against human topoisomerase IIβ was purchased from BD Transduction Laboratories (Cat. No. 611493; BD Biosciences, USA). An antibody against casein kinase 1δ (CK1δ; Cat. No. ab48031) was obtained from Abcam (Cambridge, UK), and recombinant human CK1δ protein (Cat. No. PV3665) was purchased from Invitrogen, Thermo Fisher Scientific, USA.

Dulbecco’s Modified Eagle Medium (DMEM; Cat. No. 52100-039) and Roswell Park Memorial Institute (RPMI) 1640 medium (Cat. No. 31800-089) were purchased from Gibco, Thermo Fisher Scientific, USA. Fetal bovine serum (FBS; Cat. No. 10270-106), 2.5% trypsin (Cat. No. TCL034), and penicillin–streptomycin antibiotic solution (Cat. No. A018) were also obtained from Gibco, Thermo Fisher Scientific, USA.

Mono Q anion-exchange resin (Cat. No. 17-0510-10) was obtained from GE Healthcare (Uppsala, Sweden). The ADP-Glo® Kinase Assay kit (Cat. No. V6930) was purchased from Promega (Madison, WI, USA). All other chemicals and reagents used in this study were of analytical grade.

### Methods

#### Cell culture

SupT1 and TZM-bl cells were cultured in RPMI-1640 medium (Gibco), whereas CHO_JR-FL_ and HEK293T cells (Supplementary Table S4) were cultured in DMEM/F-12 medium (Gibco), supplemented with 10% fetal bovine serum (FBS; Gibco). All cells were maintained in a humidified incubator at 37°C with 5% CO₂.

#### Isolation and culture of peripheral blood mononuclear cells (PBMCs)

Peripheral blood mononuclear cells (PBMCs) were isolated from blood collected from healthy volunteers after obtaining informed consent, in accordance with the Institutional Ethics Committee approval (UH/IEC/2021/153; dated 26 August 2021), the National Ethical Guidelines for Biomedical and Health Research Involving Human Participants issued by the Indian Council of Medical Research (ICMR), India, and the Declaration of Helsinki. PBMCs were isolated by Ficoll-Hypaque density-gradient centrifugation (Amersham Pharmacia) [48]. The isolated PBMCs were either cryopreserved in liquid nitrogen or cultured in RPMI-1640 medium supplemented with phytohaemagglutinin (PHA; 2.5 μg/mL), interleukin-2 (IL-2; 50 U/mL), penicillin (100 U/mL), and streptomycin (100 μg/mL). Following three days of activation and culture, the PBMCs were harvested for viral infection experiments.

#### Virus propagation

HIV-1 NL4-3 and isolates representing subtypes A, B, C, and D were propagated in SupT1 cells. Viral p24 antigen levels in the culture supernatants were quantified at 96 hours post-infection using a p24 ELISA kit (ABL Inc., Cat. No. 5447). The propagated viral stocks were titrated in TZM-bl cells to determine the multiplicity of infection (MOI) [49]. The recombinant HIV-1 CRF_AE isolate was propagated in PBMCs, and all subsequent experiments involving this isolate were performed using PBMCs.

#### Viral infections

All infection experiments were performed in serum-free medium at a multiplicity of infection (MOI) of 2. For protein production studies, cells were infected with HIV-1 isolates representing different subtypes. Briefly, 5 × 10⁶ cells were infected with HIV-1 at an MOI of 2 in serum-free medium. For early time-course studies, infection was terminated at 15 min, 30 min, 1 h, 2 h, and 4 h post-infection. For experiments involving longer incubation periods (>6 h post-infection), the culture medium was supplemented with 10% serum. The cells were subsequently washed and lysed using RIPA buffer containing 20 mM Tris (pH 7.5), 1 mM MgCl₂, 0.1 mM EDTA, 500 mM KCl, 1% Triton X-100, 5% glycerol, 0.1% SDS, 0.1 mM β-mercaptoethanol, 0.5 mM phenylmethylsulfonyl fluoride (PMSF), 1 μg/mL leupeptin, and 1 μg/mL pepstatin. The protein concentration of each lysate was determined, and the lysates were used for subsequent analyses.

#### Pseudovirus preparation

The HIV-1 pNL4-3ΔEnv plasmid was co-transfected with an HIV-1 subtype B envelope clone at a 1:1 ratio to generate envelope-complemented viral particles. For the generation of control virions, pNL4-3ΔEnv was co-transfected with the pVSV-G plasmid at a 1:1 ratio (Supplementary Table S2) [51]. An independent transfection with pNL4-3ΔEnv, without envelope complementation, was also performed. In a separate experiment, cells were transfected with the full-length pNL4-3 plasmid as a control. All transfections were performed in HEK293T cells using Lipofectamine 3000 according to the manufacturer’s instructions. Culture supernatants were collected 48 h post-transfection, and the viral p24 antigen concentration was quantified using an HIV-1 p24 sandwich ELISA kit (ABL Inc., Cat. No. 5447). The viral stocks were subsequently titrated using the TZM-bl assay to determine the multiplicity of infection (MOI) [49].

#### Preparation of viral lysate

Culture supernatants collected on day 4 post-infection were pooled and layered onto a 30% sucrose cushion at a 1:1 (v/v) ratio in Oak Ridge centrifuge tubes. The samples were centrifuged at 18,000 rpm for 2 h at 4°C. The supernatant was discarded, and the resulting pellet was washed once with 0.1 M Tris-buffered saline (TBS). The pellet was then resuspended in RIPA buffer to a final volume equivalent to 1/10–1/100 of the original culture supernatant volume. The resulting viral lysate was dialyzed against 0.1 M TBS for 24 h using a dialysis membrane with a 12-kDa molecular weight cut-off, with the buffer replaced every 4 h. The dialyzed sample was subsequently concentrated, and the protein concentration was determined using the bicinchoninic acid (BCA) assay. The final sample was stored at −80°C until further use.

#### Purification of TopoIIβK from the virus

TopoIIβK was purified from the dialyzed viral lysate by anion-exchange chromatography using a Mono Q column (GE Healthcare) at 4°C. The column was packed with Mono Q resin to a bed height of 1–1.5 cm and equilibrated with 20 bed volumes of Tris-buffered saline (TBS). The viral lysate was subsequently loaded onto the column, followed by washing and sequential elution using buffers containing increasing NaCl concentrations (100, 200, 400, and 800 mM). Following elution, the column was washed with TBS and stored at 4°C. The protein concentration in the collected fractions was determined, and the purified protein was stored at −80°C until further use.

#### Purification of TopoIIβK from gp120-expressing cells

##### CHO_JR-FL_ cells

CHO_JR-FL_ cells were cultured in DMEM/F-12 medium. As these cells secrete gp160 into the culture medium, the culture supernatant was collected for the purification of TopoIIβK by Mono Q anion-exchange chromatography, as described above.

##### Transfection of gp120 clones in HEK293T cells

HEK293T cells (2.5 × 10⁶) were transfected with 2 μg of plasmid DNA using Lipofectamine 3000 (Gibco), according to the manufacturer’s instructions. At 36 h post-transfection, the cells were harvested and lysed using RIPA buffer. TopoIIβK was subsequently purified from the cell lysate by Mono Q anion-exchange chromatography, as described above.

#### MALDI ToF/ToF analysis

The procedure was performed as described by Kondapi et al. [11]. Briefly, TopoIIβK was purified by Mono Q anion-exchange chromatography, and the bound protein was step-eluted using increasing concentrations of NaCl in 20 mM Tris-HCl buffer (pH 8.6). The fraction eluted at 800 mM NaCl was collected for further analysis. The purified protein was digested with trypsin, and the resulting peptides were analyzed by MALDI-TOF/TOF. The detected ions were annotated with their monoisotopic masses and the corresponding amino acid positions in the protein sequence. The resulting mass spectrum was searched against the NCBI protein database using the Mascot search engine to identify proteins matching the experimental mass spectral data.

#### Western blot analysis

Western blotting was performed as previously described by Lokeswara et al. [7]. Briefly, 100 µg of total protein was separated by 10% SDS–polyacrylamide gel electrophoresis (SDS-PAGE; Thermo Fisher Scientific) and subsequently transferred onto a 0.45-µm polyvinylidene difluoride (PVDF) membrane (Millipore). The membrane was incubated with a mouse anti-TopoIIβK primary antibody, followed by incubation with a horseradish peroxidase (HRP)- conjugated goat anti-mouse IgG secondary antibody (Cell Signaling Technology). Immunoreactive bands were detected using an enhanced chemiluminescence (ECL) reagent, and the resulting chemiluminescent signals were captured using a ChemiDoc imaging system (Bio-Rad).

#### Immunofluorescence to check the production of Kinase in lysosomes

TZM-bl cells were incubated with LysoTracker (Thermo Fisher Scientific) for 1 h at 37°C and subsequently infected with HIV-1 NL4-3 at a multiplicity of infection (MOI) of 2. At the indicated post-infection time points (15, 30, 45, and 60 min), the cells were fixed with 4% formaldehyde for 15 min at room temperature and washed twice with phosphate-buffered saline (PBS). The cells were then permeabilized with 0.1% Triton X-100 for 10 min, washed with PBS, and incubated overnight at 4°C with a mouse anti-TopoIIβK primary antibody. After two PBS washes, the cells were incubated with an Alexa Fluor 488-conjugated goat anti-mouse secondary antibody (Invitrogen) for 2 h at room temperature in the dark. The nuclei were counterstained with Hoechst dye, and fluorescence images were acquired using a Carl Zeiss confocal laser-scanning microscope.

#### Effect of cysteine protease inhibitor on TopoIIβK production

gp120 transfection studies: TZM-bl cells (2.5 × 10⁶) were preincubated with 10 μM of the cysteine protease inhibitor E64d (Tokyo Chemical Industry) or the cathepsin L inhibitor Z-FY-CHO (Sigma-Aldrich). The cells were subsequently transfected with the HIV-1 subtype B envelope plasmid pTRO11 (NIH, USA) using Lipofectamine 3000 (Invitrogen), according to the manufacturer’s instructions. Cell pellets were collected at 12, 24, and 48 h post-transfection for further analysis.

E64d treatment during HIV-1 infection: SupT1 cells (4 × 10⁶) were preincubated with 10 μM E64d and subsequently infected with HIV-1 NL4-3 at a multiplicity of infection (MOI) of 2. Cells were harvested at 15 min, 30 min, 1, 2, 4, 12, 24, 48, 72, and 96 h post-infection (p.i.). The cell lysates were analyzed by Western blotting following separation by 10% SDS–polyacrylamide gel electrophoresis (SDS-PAGE).

Z-FY-CHO treatment during HIV-1 infection: TZM-bl cells (2.5 × 10⁶) were preincubated with 10 μM Z-FY-CHO (Sigma-Aldrich) and subsequently infected with HIV-1 NL4-3 at an MOI of 2. Cells were harvested at 15 min, 30 min, 1, 2, 4, and 12 h p.i. TopoIIβK expression was subsequently assessed by Western blotting after separation of cell lysates on 10% SDS-PAGE gels.

#### ADP-Glo® kinase assay

The kinase activity of TopoIIβK was measured using the ADP-Glo® Kinase Assay (Promega, Madison, WI, USA), according to the manufacturer’s instructions. The TopoIIβ substrate was prepared as described previously [11]. Before initiating the kinase reaction, the substrate was washed twice with kinase buffer containing 50 mM Tris-HCl (pH 7.5), 5% glycerol, 10 mM MgCl₂, and 5 mM dithiothreitol (DTT) to remove residual phosphatase activity.

The kinase reaction was performed by incubating purified TopoIIβK with TopoIIβ in kinase buffer supplemented with 100 μM ATP at 37°C for 30 min. Each 50-μL reaction contained 1 μM TopoIIβ as the substrate and 50 nM TopoIIβK purified from different HIV-1 subtypes. Purified casein kinase 1δ (CK1δ) was used as a positive control.

Following the kinase reaction, 50 μL of ADP-Glo® Reagent was added to each reaction mixture, then incubated at room temperature for 40 min to terminate the kinase reaction and deplete any remaining ATP. Subsequently, 100 μL of Kinase Detection Reagent was added, and the samples were incubated at room temperature for 30 min to convert ADP to ATP and generate a luminescent signal proportional to the amount of ADP produced. Luminescence was measured using a microplate luminometer with an integration time of 0.5–1 s per well. The measured luminescence, expressed as relative luminescence units (RLUs), was converted to the amount of ADP produced (pmol) using an ADP/ATP standard curve. All experiments were performed using at least three independent biological replicates, and the data are presented as the mean ± standard deviation (SD).

#### Cloning, expression and purification of recombinant TopoIIβK (precTopoIIβK)

Based on the immunoreactivity of TopoIIβK, the N-terminal region of HIV-1 gp120 was PCR-amplified from the pNL4-3 template and cloned into the pEGFP-C1 vector using the EcoRI and SalI restriction sites. The resulting construct was verified by restriction digestion and Sanger sequencing. HEK293T cells were cultured in DMEM supplemented with 10% fetal bovine serum (FBS) at 37 °C in a humidified atmosphere containing 5% CO₂. The cells were transfected with the recombinant construct using Lipofectamine 3000, according to the manufacturer’s instructions. At 36 h post-transfection, the cells were harvested and lysed in RIPA buffer supplemented with protease inhibitors.

The clarified cell lysate was subjected to anion-exchange chromatography using a Mono Q column (GE Healthcare) equilibrated with 20 bed volumes of Tris-buffered saline (TBS) at 4 °C. Bound proteins were eluted stepwise using TBS containing 100, 200, 400, and 800 mM NaCl. The collected fractions were analyzed for the presence of the target protein, and the relevant fractions were pooled. Protein concentration was determined using the bicinchoninic acid (BCA) assay, and the purified protein was stored at −80 °C. Native TopoIIβK was purified from concentrated HIV-1 virion lysates using the same Mono Q chromatography procedure.

#### Comparative analysis of native and recombinant TopoIIβK

Kinase activity of native TopoIIβK and precTopoIIβK was determined using the ADP-Glo® Kinase Assay (Promega, Madison, WI, USA) according to the manufacturer’s instructions. The substrate TopoIIβ was prepared as described earlier [11] and washed twice with kinase buffer (50 mM Tris pH 7.5, 5% glycerol, 10 mM MgCl₂ and 5 mM DTT) to remove residual phosphatase before the kinase reaction was started.

Kinase reactions (50 µL) were set up by incubating purified native TopoIIβK or precTopoIIβK with 1 µM TopoIIβ in kinase buffer containing 100 µM ATP at 37 °C for 30 min. To assess dose dependence, enzyme concentrations of 10, 20, 25, 50, and 100 nM were used. An equal volume (50 µL) of ADP-Glo® Reagent was then added, and the samples were incubated for 40 min at room temperature to terminate the reaction and deplete unconsumed ATP. Kinase Detection Reagent (100 µL) was added, and the samples were incubated for a further 30 min at room temperature to convert ADP to ATP and generate a luminescent signal proportional to the ADP produced. Luminescence was measured in a microplate luminometer with an integration time of 0.5–1 s per well. Relative luminescence units (RLU) were converted to pmol ADP using an ADP/ATP standard curve, and the background signal was subtracted.

#### Inhibition of TopoIIβK activity

To assess inhibitor sensitivity, kinase reactions were performed using 1 µM TopoIIβ and 100 nM native TopoIIβK or precTopoIIβK in kinase buffer. UHAKKM-12, a dicoumarin-based inhibitor of TopoIIβK, was added at final concentrations of 100, 300, 600, 700, and 800 nM. The kinase reaction was initiated by adding ATP to a final concentration of 100 µM, followed by incubation at 37 °C for 30 min. The reactions were subsequently processed using the ADP-Glo® Kinase Assay, as described above. UHAKKM-12 exhibited an IC₅₀ of 85.26 nM for native TopoIIβK and 294.86 nM for precTopoIIβK, under the conditions tested.

## Results

### Kinetics of TopoIIβK formation during HIV-1 infection

The kinetics of TopoIIβK formation during HIV-1 post-infection (p.i.) were analyzed using subtype 99KE_KNH1088 (Subtype A), 90US_873 (Subtype B), 94IN_20635-4 (Subtype C), and 98UG_57128 (Subtype D) viruses in Sup T1 cells (Fig. 1). The results show that in subtype A, TopoIIβK appears as two bands with consistent production from 0.5 h post-infection (p.i.) to 4 h p.i. (Fig. 1A); while in human PBMCs, TopoIIβK was observed from 0.25 h, decreasing gradually until 2 h (Fig. 1B). In subtype B and C infections, two bands were observed, with the higher-molecular-weight band (72 kDa) corresponding to TopoIIβK. In subtype B, TopoIIβK appeared at 0.25 h p.i., then decreased at subsequent time points, but remained significant until 4 h. In the case of subtype C infection, TopoIIβK was observed from 0.25 h to 1 h, followed by a sharp decrease from 2 h. Subtype D has been shown to produce TopoIIβK from 0.25 h, reaching significant levels at 0.5 h, followed by a gradual decrease at later p.i. Further, for the infection using HIV-1 recombinant 96TH_M02138 (CRF_AE), TopoIIβK appeared as a single band at 0.25 h and then decreased at subsequent time points. Overall, we observe that TopoIIβK appears at a very early time point, 0.25 h post-infection, followed by changes in its level across the virus subtypes, with altered kinetics that may correlate with viral diversity and different host cell environments, namely SupT1 cells and PBMCs.

**Figure 1.**
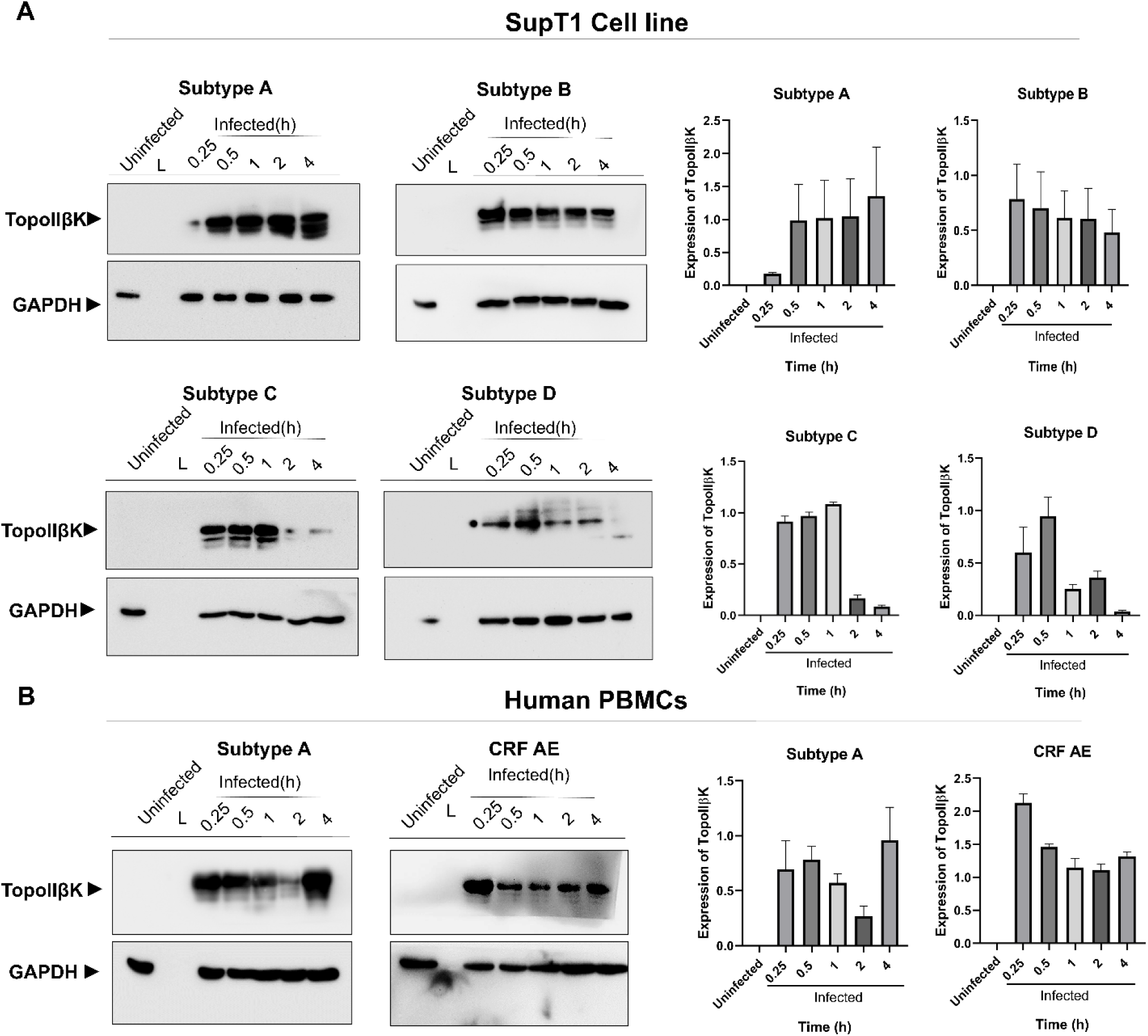
The production of Topoisomerase II β kinase is conserved among HIV-1 subtypes. The dynamics of TopoIIβK production in the early stages of infection (0.25 to 4 hours p.i.) were assessed using HIV-1 subtypes at different time points. The level of TopoIIβK was analyzed by Western blot with a mouse anti-TopoIIβK antibody. **(A)** HIV-1 subtypes A-99KE_KNH1088, B-90US_873, C-94IN_20635-4, and D-98UG_5712 in SupT1 cell line; **(B)** HIV-1 subtype (A-99KE_KNH1088) and CRF AE (96TH_M02138) in human PBMCs. Graphical representation of densitometric analysis of TopoIIβK protein levels normalized against their respective GAPDH expression levels. Data represent mean ± SD from three independent experiments performed in triplicate (n = 3).

### TopoIIβK produced in HIV-1 envelope-expressing cells

When we analyzed the cell supernatant from CHO_JR-FL_ for TopoIIβK production (Fig. 2A), we detected a significant amount of a 72-kDa protein in the culture medium, which was purified by Mono-Q ion-exchange chromatography and confirmed as TopoIIβK (Fig. 2B).

**Figure 2.**
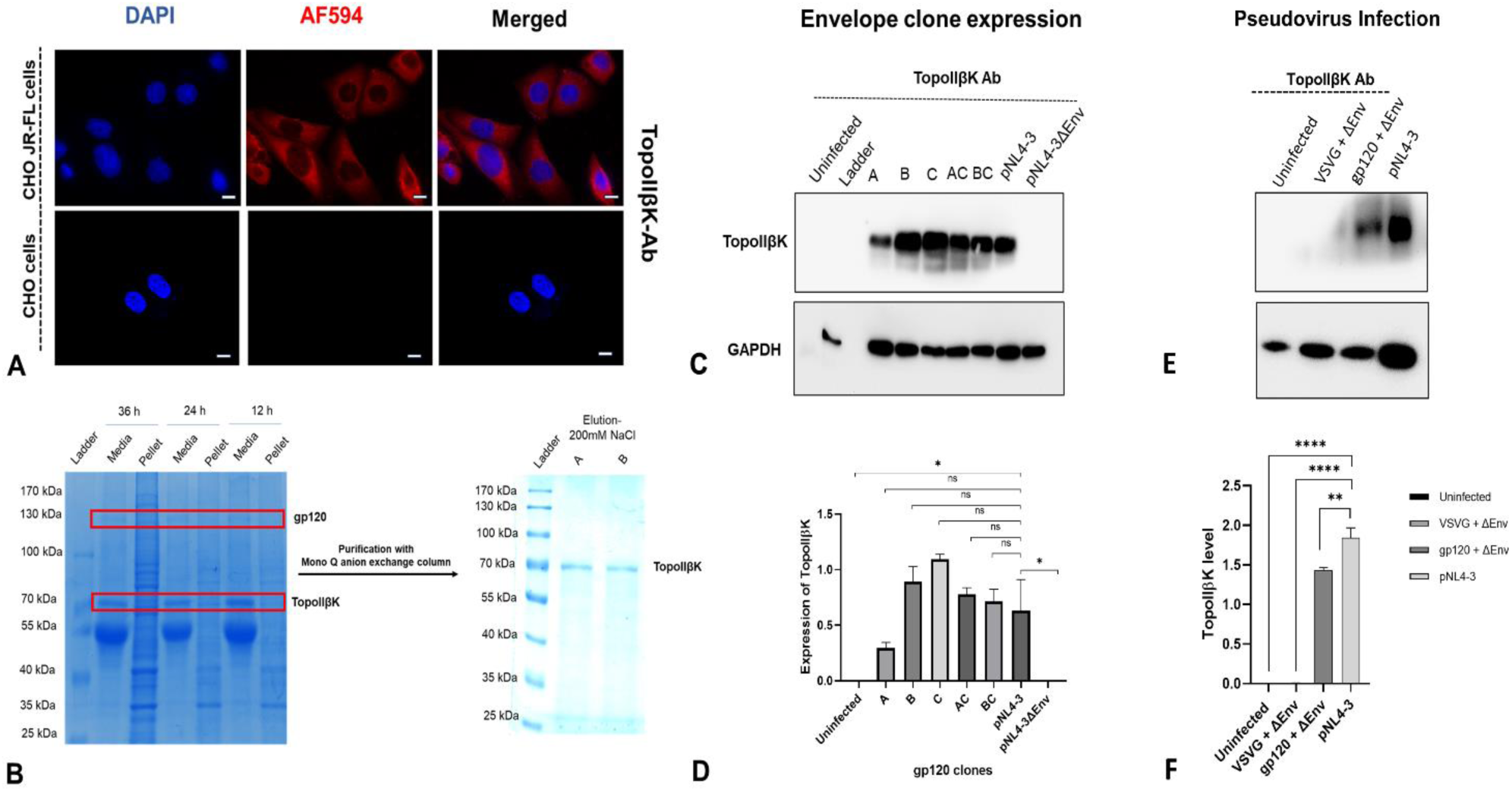
Topoisomerase II β kinase is produced in HIV-1 envelope-expressing cells. The production of TopoIIβK was analyzed in envelope-expressing cells, as indicated. **(A)** Immunofluorescence studies of envelope-expressing CHO_JR-FL_ cells using an anti-TopoIIβK antibody with an Alexa Fluor 594-tagged secondary anti-mouse antibody. Images were captured using a Carl Zeiss fluorescence microscope Axiocam 3.0. Scale bar represents a length of 10μm; **(B)** Purification of TopoIIβK from CHO_JR-FL,_ as assessed by SDS-PAGE, the results show TopoIIβK is secreted in these cells, which can be purified by Mono-Q anion exchange chromatography; **(C)** HEK293T cells were transfected using gp120 plasmids from subtype A, B, C, AC, and BC viruses viz. p398F1env, pTRO11env, pCE1176env, p246F3env, and pBJOX2000. The transfected cells were harvested after 36 hours, and TopoIIβK production was analyzed by Western blotting using a mouse anti-TopoIIβK antibody. Also, lanes 8 and 9 show the production of TopoIIβK in cells transfected with pNL4-3 complete and pNL4-3 ΔEnv clones; **(D)** represents the densitometric analysis of TopoIIβK protein levels in (C), normalized against their respective GAPDH expression level (mean ± SD, n = 3). **(E)** TZM-bl cells were infected for one hour with (i) none; (ii) pseudovirus pNL4-3ΔEnv complemented with pVSVG; (iii) pseudovirus pNL4-3ΔEnv complemented with subtype B pTRO11; (iv) complete infectious clone pNL4-3. The infected cells were harvested, and TopoIIβK production was analyzed by Western blot with a mouse anti-TopoIIβK antibody. **(F)** represents the densitometric analysis of TopoIIβK protein levels in (E), normalized to their respective GAPDH expression levels (mean ± SD, n = 3). Statistical significance was determined by a paired t-test with Welch’s correction; \**p* < 0.05, \*\**p* < 0.01, \*\*\*\**p* < 0.0001.

To independently verify, HIV-1 envelope clones, viz. p398F1 (Subtype A), pTRO11 (Subtype B), pCE1176 (Subtype C), p246F3 (Subtype AC recombinant), and pBJOX2000 (CRF_BC) were transfected into HEK293T cells. The expressed protein at 36 h was analyzed by Western blot using a mouse anti-TopoIIβK antibody; the results in Fig. 2C and 2D show TopoIIβK production in transfected cells, suggesting that TopoIIβK is expressed from the envelope of all four subtypes as well as from recombinants. Although there is significant formation of TopoIIβK, protein levels vary among envelope clones derived from different subtypes; this may reflect intrinsic sequence variation among these viruses.

To confirm whether TopoIIβK is indeed derived from the envelope, we transfected the pNL4-3 infectious plasmid and the pNL4-3ΔEnv plasmid in HEK293T cells. The protein was analyzed by Western blot; the results in Fig. 2C and 2D, lanes 8 and 9, show that TopoIIβK is produced in cells expressing the complete infectious pNL4-3, whereas it is not produced in the envelope-deleted clone (pNL4-3ΔEnv). Further, to reconfirm, we have complemented the pNL4-3ΔEnv plasmid with HIV-1 gp120 and, in another experiment, with pVSV-G. The results in Fig. 2E and 2F show a distinct 72-kDa TopoIIβK band in TZM-bl cells infected with the gp120-complemented pseudovirus, similar to that observed with the HIV-1 NL4-3 positive control. In contrast, no detectable TopoIIβK signal was observed in pNL4-3ΔEnv-infected cells complemented with pVSV-G, i.e., a pseudovirus expressing VSV-G. The results indicate that VSV-G-mediated entry is insufficient to produce TopoIIβK, as VSV-G may not contain TopoIIβK. These results further point out the importance of gp120 in the production of TopoIIβK. These results clearly demonstrate that TopoIIβK originates from gp120.

### TopoIIβK originates from the N-terminus of the HIV-1 envelope and is conserved among subtypes

TopoIIβK was purified using Mono-Q ion-exchange chromatography (Fig. 3A). TopoIIβK was trypsin-digested and analyzed using MALDI-ToF/ToF (Fig. 3B). Each ion generated in the analysis has been labeled by both the monoisotopic mass and the amino acid position in the protein sequence. Mascot analysis of peptides identified a single peptide homologous to the HIV-1 envelope glycoprotein, with a high likelihood score of 82 (Supporting data Fig. S1). Furthermore, we compared this peptide sequence with the envelope glycoproteins of subtypes A, B, C, D, AE, and pNL4-3 (Supporting data Fig. S1B). The multiple sequence alignment clearly showed a significant match between the peptide and the envelope glycoprotein of all subtypes around the V3 region. Furthermore, the analysis showed that peptides generated from MS/MS data covered 60% of the HIV-1 envelope glycoprotein sequence (Fig. 3C).

**Figure 3:**
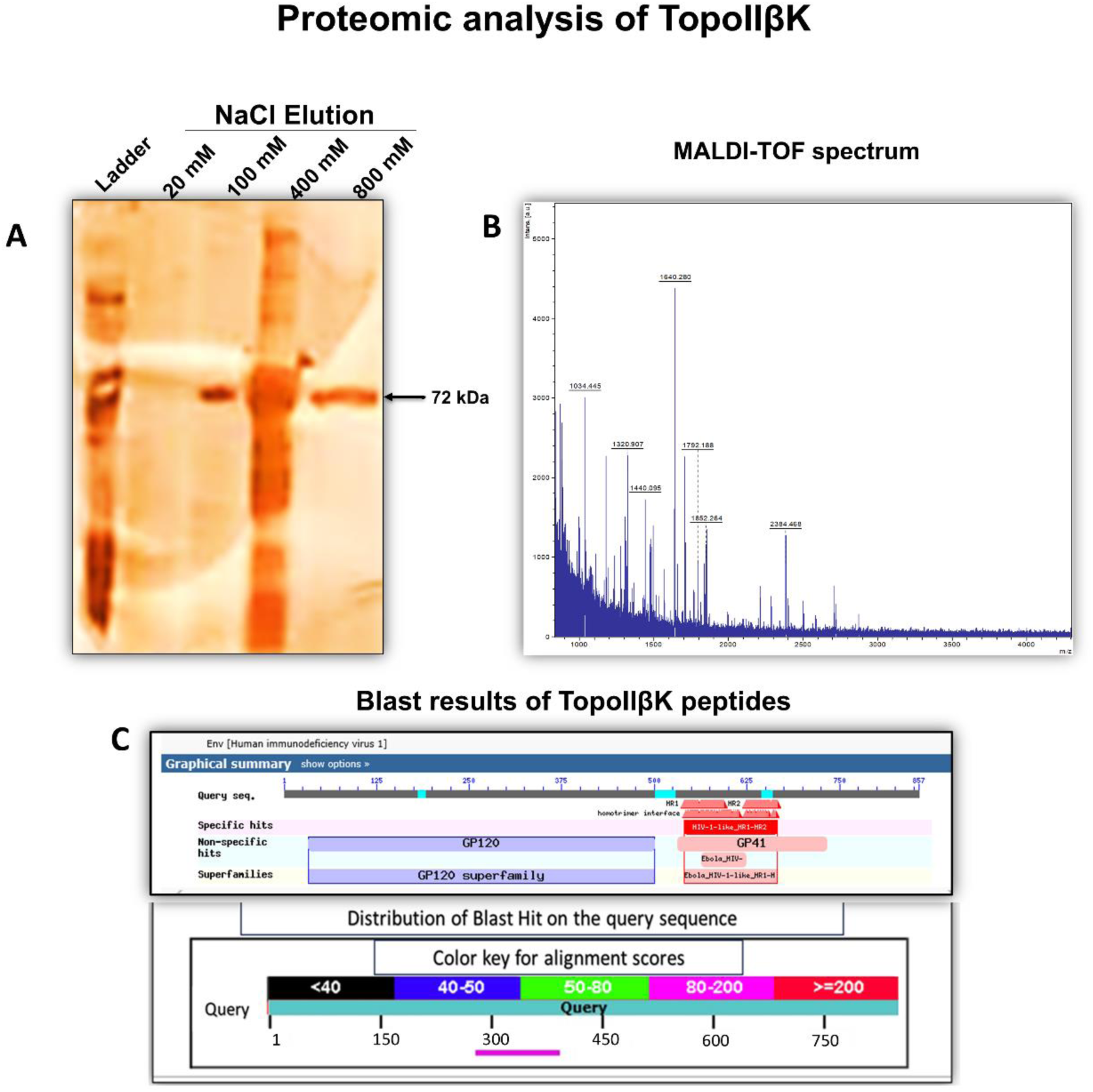
Proteomic analysis of TopoIIβK. **(A)** Silver-stained gel showing eluted protein through the Mono-Q column chromatography using different concentrations of NaCl. The arrow indicates a 72 kDa TopoIIβK band at 100 mM NaCl and 800 mM NaCl. **(B)** The TopoIIβK protein in the 800 mM NaCl eluate was excised and sent for MALDI-TOF/TOF analysis. The figure shows the mass spectrum generated from MALDI-TOF analysis, and **(C)** Peptides obtained from the MS/MS data of MALDI-TOF analysis were further aligned with the gp160 protein sequence of HIV-1, showing that out of 857 amino acids, the peptides covered around 287 to 390 amino acids, confirming that the 72 kDa TopoIIβK protein may be derived from the HIV-1 gp120 region of the envelope.

To explore the common attributes of TopoIIβK and HIV-1 gp120, the purified viral TopoIIβK was assessed by Western blot using a panel of mouse anti-gp120 monoclonal antibodies (Fig. 4A), as indicated [24,25,26]. Among the antibodies used, 2909, CH58, and ID6 recognized TopoIIβK (Fig. 4B), suggesting that TopoIIβK may be derived from the N-terminal portion of gp120.

**Figure 4.**
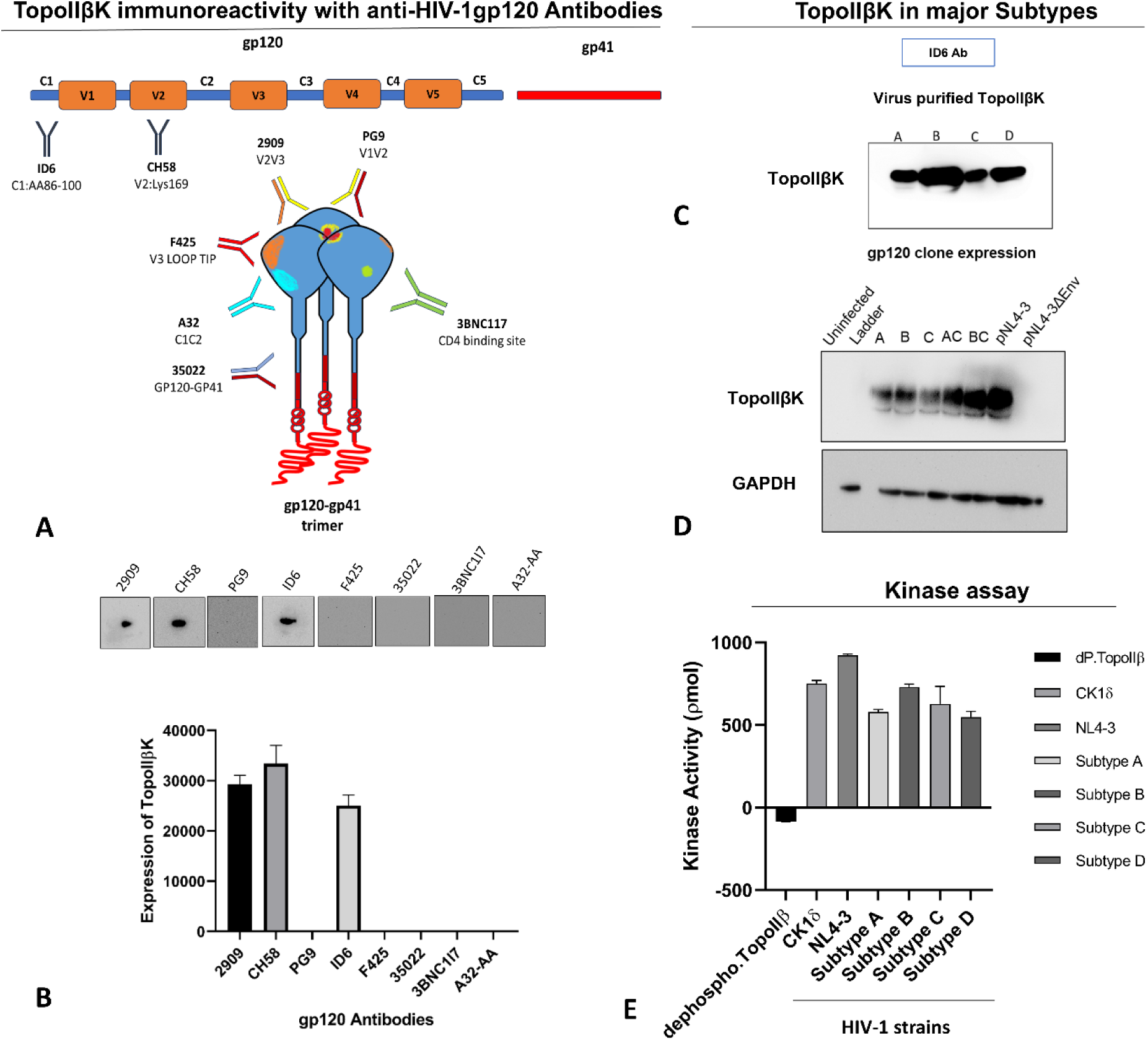
TopoIIβK is derived from the N-terminal portion of gp120. Immunoreactivity of TopoIIβK against mouse-HIV-1 anti-gp120 monoclonal antibodies: **(A)** Graphical representation of binding specificity of HIV-1 anti-gp120 monoclonal antibodies used in the analysis; **(B)** Purified TopoIIβK from HIV-1NL4-3 was analyzed by Western blotting using the indicated antibodies in panel (A). The results show that TopoIIβK is immunoreactive to ID6 (recognizes 86-100 amino acids of gp120), CH58 (V2 epitope), and 2909 (V2-V3 quaternary epitope), suggesting TopoIIβK is derived from the N-terminal of gp120; **(C)** Shows immunoreactivity of TopoIIβK purified from HIV-1 subtypes A-99KE_KNH1088, B-90US_873, C-94IN_20635-4, and D-98UG_57128 viruses. The purified TopoIIβK was Western-blotted with ID6 antibody; the results show that TopoIIβK from all subtypes is recognized by ID6, suggesting that TopoIIβK is derived from the N-terminus of gp120 and is conserved. **(D)** Shows immunoreactivity of TopoIIβK purified from HIV-1 gp120 clones derived from subtype A, B, C, AC, and BC viruses, viz. p398F1env, pTRO11env, pCE1176env, p246F3env, and pBJOX2000. The purified TopoIIβK was analyzed by Western blotting with ID6 antibody; the results also show that TopoIIβK produced in gp120-expressing cells is recognized by ID6, suggesting that TopoIIβK is derived from the N-terminus of gp120 and is conserved among subtypes. **(E)** TopoIIβK kinase activity was measured using the ADP-Glo kinase assay for TopoIIβK isolated from HIV-1 strain NL4-3 and subtypes A, B, C, and D. TopoIIβ was used as a negative control, and CK1δ was used as a positive control. The kinase activity is derived from the ADP-Glo standard curve (Supporting data Fig. S7). The data are represented as pmol of ADP generated, mean ± SD, from three independent experiments performed in triplicate (n = 3).

To determine whether the mouse anti-TopoIIβK antibody recognizes the host kinase CK1δ [50], immunoblot analysis was performed using purified TopoIIβK, lysates from HIV-1-infected and uninfected cells, and purified CK1δ. Immunoblotting with the anti-CK1δ antibody detected CK1δ in both infected and uninfected cell lysates, with the strongest signal observed for the purified CK1δ control, whereas no detectable signal was observed for purified TopoIIβK (Supporting information Fig. S2). In contrast, the anti-TopoIIβK antibody strongly recognized purified TopoIIβK and detected the kinase in HIV-1-infected cell lysates but showed no detectable reactivity with purified CK1δ or uninfected cell lysates.

The purified TopoIIβK from all subtypes (Supporting information Fig. S3) was analyzed by Western blotting using a mouse anti-gp120 (N-terminal 86-100 aa) antibody [27,28,29]; the results in Fig. 4C show that TopoIIβK is cross-reactive with the N-terminal gp120 antibody, ID6. These results suggest that TopoIIβK is derived from the N-terminal region of gp120 and is conserved across a broad spectrum of virus strains.

Furthermore, HEK293T cells were transfected with p398F1 (Subtype A), pTRO11 (Subtype B), pCE1176 (Subtype C), p246F3 (Subtype AC recombinant), and pBJOX2000 (CRF_BC), and lysates were prepared 36 hours post-transfection. The lysates were analyzed for TopoIIβK by Western blot using the mouse anti-gp120 monoclonal antibody ID6; the results in Fig. 4D (Supporting Information Fig. S4) show that TopoIIβK produced from gp120 clones of different subtypes was cross-reactive with the ID6 antibody, which recognizes the N-terminal portion of gp120, suggesting that TopoIIβK is derived from the N-terminus of gp120.

We have purified TopoIIβK from different viral subtypes (see Supporting Information, Fig. S3). The biochemical activity of purified TopoIIβK in terms of phosphorylation of the substrate Topoisomerase II β was analyzed using the ADP-Glo® Kinase Assay. The results shown in Fig. 4E indicate that TopoIIβK exhibits kinase activity across all subtypes. The negative control, TopoIIβ substrate alone, showed (negative) 86.09 pmol, confirming the negligible background phosphorylation in the absence of an active kinase; also, there is no evidence of co-purified kinase with TopoIIβ. In contrast, the CK1δ [50] positive control displayed robust kinase activity (752.46 pmol), validating the assay system.

Among the HIV-1 strains tested, the TopoIIβK purified from laboratory-adapted NL4-3 strain exhibited the highest kinase activity (922.76 pmol), exceeding that of the CK1δ control. The TopoIIβK purified from primary HIV-1 subtypes examined also demonstrated substantial kinase activity, albeit with some variation: Subtype A (579.53 pmol), Subtype B (730.72 pmol), Subtype C (627.67 pmol), and Subtype D (547.73 pmol). TopoIIβK purified from all HIV-1 strains efficiently phosphorylated TopoIIβ *in vitro*, confirming that TopoIIβ is a substrate of TopoIIβK (Fig. 4E, Supporting Information Fig. S5). These findings indicate that TopoIIβK-mediated phosphorylation of TopoIIβ is conserved across HIV-1 strains.

### Recombinant partial TopoIIβK from N-terminal gp120 exhibits kinase activity

Based on the immunoreactivity of TopoIIβK, we have cloned the N-terminal region of gp120 (precTopoIIβK) into the pEGFP-C1 vector and expressed it in HEK293T cells. The expressed protein was purified using Mono Q affinity chromatography. The results presented in Fig. 5A show that precTopoIIβK was recognized by the anti-TopoIIβK antibody. The kinase activity of precTopoIIβK was assessed by analyzing the phosphorylation of 180 kDa TopoIIβK catalyzed by precTopoIIβK using the ADP-Glo assay. The results presented in Fig. 5B show that precTopoIIβK possesses significant phosphorylation activity comparable to that of virus-purified TopoIIβK against host 180 kDa TopoIIβ substrate. Furthermore, both precTopoIIβK and native TopoIIβK show dose-dependent sensitivity to UHAKKM-12 (a dicoumarin inhibitor of TopoIIβK) with IC50 of 85.26 nM for native TopoIIβK and 294.86 nM for precTopoIIβK (Fig. 5C).

**Figure 5.**
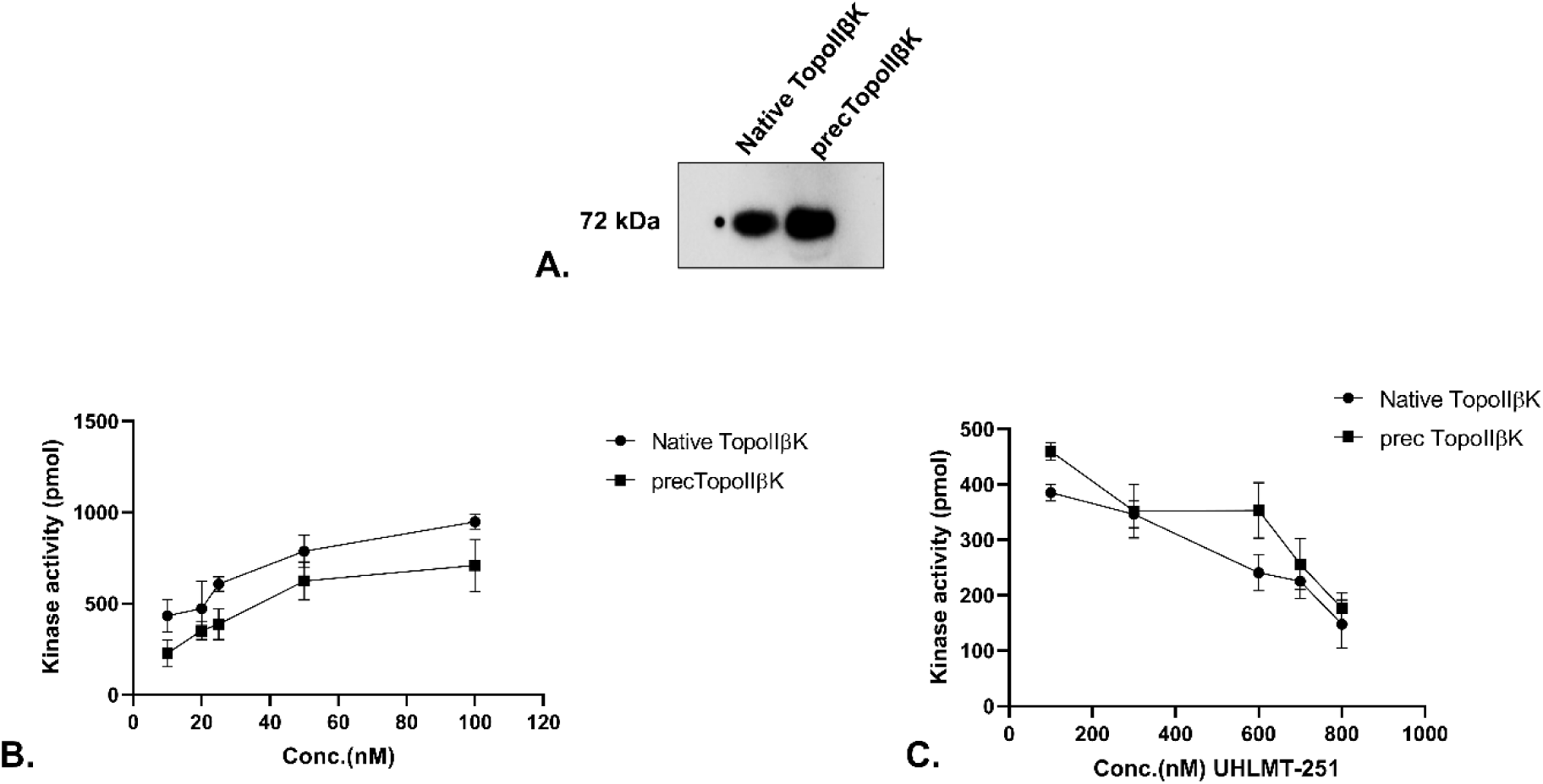
Catalytic activity of native and recombinant TopoIIβK and their sensitivity to UHAKKM-12. (A) Immunoblot of native TopoIIβK (purified from HIV-1 virions) and recombinant TopoIIβK (precTopoIIβK) probed with anti-TopoIIβK antibody, showing a single band at ∼72 kDa. (B) Dose-dependent kinase activity of native (circles) and recombinant (squares) TopoIIβK (10-100 nM), measured by the ADP-Glo assay with TopoIIβ as the substrate and expressed as pmol of ADP produced. (C) Effect of increasing concentrations of UHAKKM-12 (100-800 nM) on the kinase activity of native (circles) and recombinant (squares) TopoIIβK. Data are the mean ± SD of 3 independent experiments, each performed in triplicate.

### Localization of TopoIIβK in endolysosomes in infected cells

To examine the subcellular localization of TopoIIβK during early HIV-1 infection, TZM-bl cells were labeled with LysoTracker, infected with HIV-1 NL4-3 (MOI 2), and analyzed by confocal microscopy at early post-infection time points. The results presented in Fig. 6 show that TopoIIβK exhibited prominent co-localization with LysoTracker-positive endolysosomal compartments at 15 min post-infection (Fig. 6, Panel A). The transient enrichment of TopoIIβK within endolysosomal compartments at 15 min post-infection suggests that TopoIIβK is formed in endolysosomes of infected cells as early as 15 min; the protein overlaid on lysosomes appears near the cell membrane, while at later time points, TopoIIβK becomes more prominent and moves away from the endolysosome (Fig. 6, Panel B-D). These results indicate that TopoIIβK may be generated in the endolysosome and released into the cytosol, where it participates in HIV-1 replication. To determine whether the observed localization is due to cross-reactivity of anti-TopoIIβK with gp120, we conducted western blot analysis of gp120 and TopoIIβK using an anti-TopoIIβK antibody. The results in Supporting Information Figures 8A and 8B show that gp120 does not cross-react with the anti-TopoIIβK antibody and that it detects a 72 kDa protein. To determine the reason, we analyzed the secondary structure of TopoIIβK and gp120 using CD spectroscopy; the results in Supporting Information Fig. 8C show a significant difference in secondary structure between these proteins. Whether such a difference in their structure contributes to the differential recognition by the anti-TopoIIβK antibody requires further investigation. The results of these experiments rule out the possibility that the observed kinetics of TopoIIβK localization in endosomes are due to cross-reactivity between anti-TopoIIβK and gp120.

**Figure 6.**
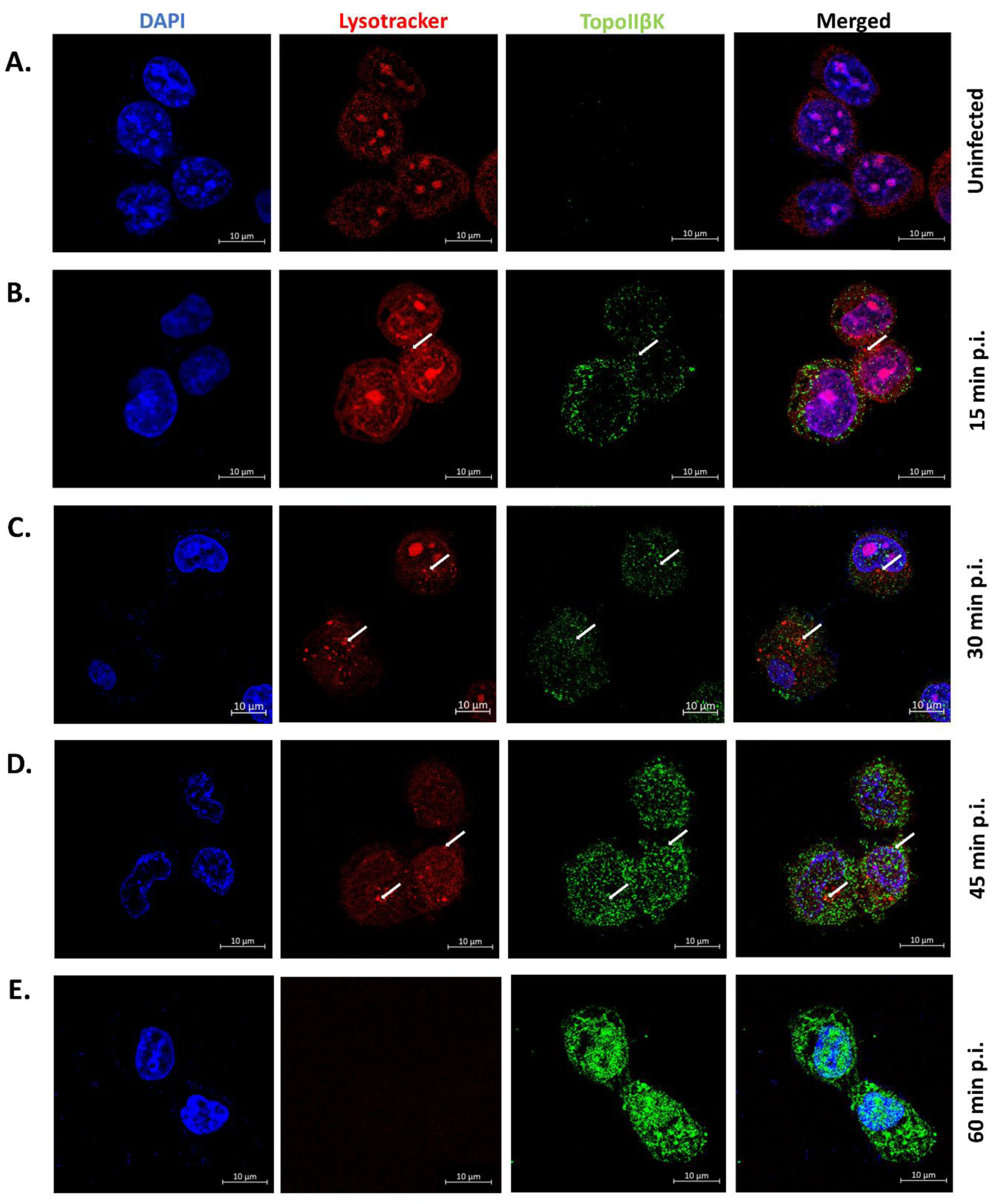
Dynamic redistribution of TopoIIβK during early HIV-1 infection. TZM-bl cells were labeled with LysoTracker and infected with HIV-1 NL4-3 (MOI, 2); the infection was stopped at the indicated time points, and cells were fixed with formaldehyde. These cells were observed under a confocal microscope. **(A)** At 15 min postinfection, TopoIIβK exhibits pronounced colocalization with LysoTracker-positive endolysosomal compartments, predominantly at the cell periphery. **(B–D)** At later early postinfection time points, TopoIIβK fluorescence becomes more intense and progressively dissociates from LysoTracker-positive compartments, with signal detected away from endolysosomes, consistent with cytosolic redistribution.

### TopoIIβK may be a product of cysteine protease-mediated cleavage of gp120

HIV-1 gp120 undergoes proteolytic cleavage by various cellular proteases and endopeptidases [32,33,34]. Among these, lysosomal cysteine proteases, such as cathepsins, undergo cellular repositioning and secretion, and modulate MHC-II peptide binding and presentation during infection [31]. Cathepsin L and B correspond to the largest share in the lysosomal cathepsin pool [30]. Thus, we have studied TopoIIβK production during HIV-1 infection and in envelope-expressing cells in experiments conducted in the presence of the cysteine protease inhibitor E64d. Based on the cytotoxicity of E64d in SupT1 and TZM-bl cells (GI_50_=>40 µM), we selected a 10 µM concentration for these experiments (Supporting data Fig. S6).

When TZM-bl cells were pre-incubated with E64d for 1 h, followed by transfection with a subtype B (pTRO11) envelope clone, TopoIIβK was analyzed at 12 h, 24 h, and 48 h post-transfection. The results in Fig. 7A show a significant decrease in TopoIIβK levels, thereby abolishing TopoIIβK formation at 48 h. These results suggest that cysteine proteases may be involved in the processing of envelope formation in TopoIIβK in envelope-transfected cells.

**Figure 7.**
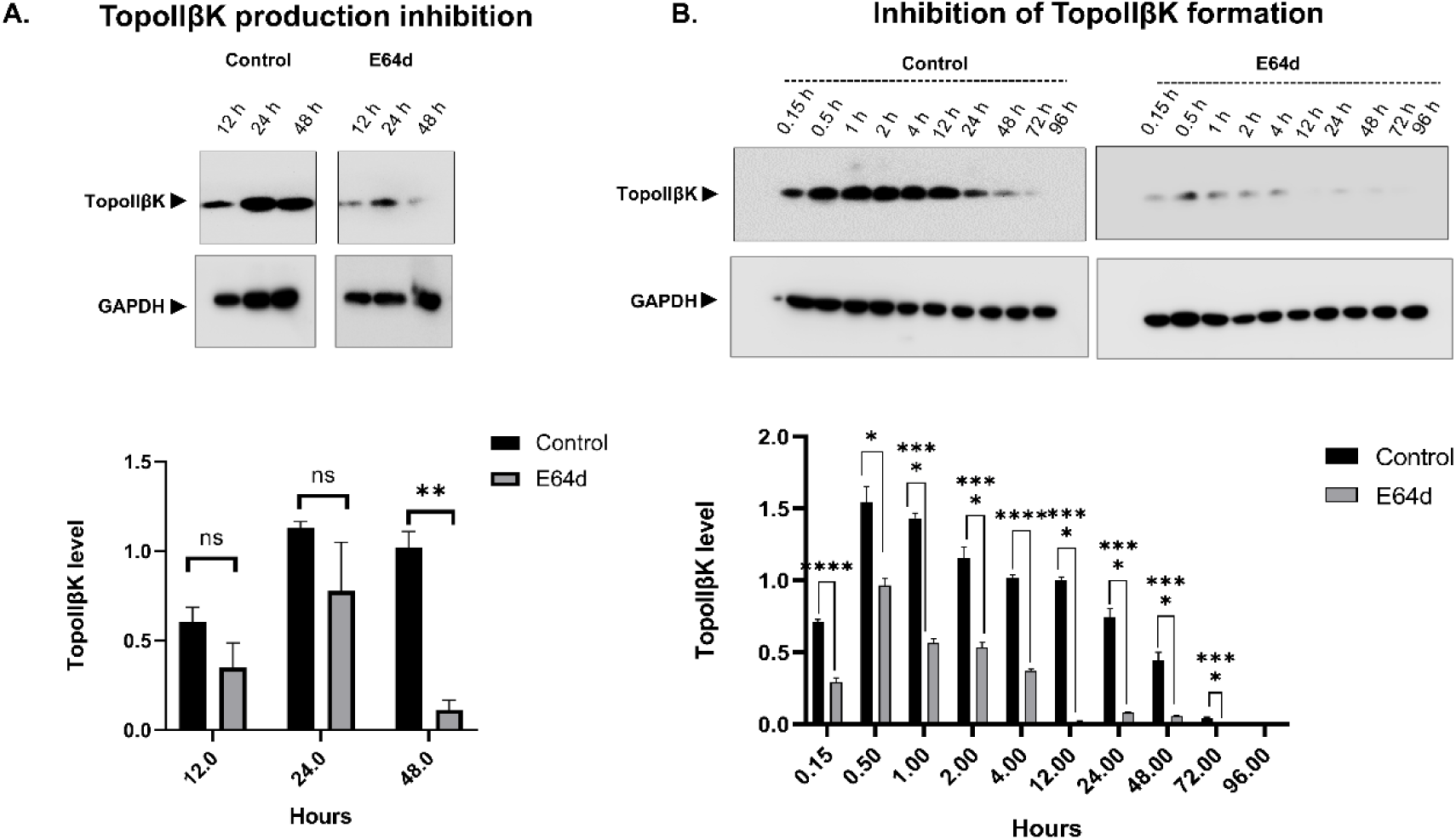
Cysteine protease inhibitor E64d inhibits TopoIIβK production. **(A)** TZM-bl cells were pre-incubated for 1 hour with 10 µM cysteine protease inhibitor E64d and then transfected with subtype B gp120 (pTRO11) plasmid. The cells were harvested at the indicated time points, and the production of TopoIIβK was analyzed by Western blot using a mouse anti-TopoIIβK antibody. Transfection in the absence of E64d is included as a negative control. The results show a significant reduction of TopoIIβK production in E64d-treated cells. **(B)** Sup T1 cells were pre-incubated with 10 µM E64d for 1 hour. These cells were infected with HIV-1 NL4-3 (2 MOI). Infection was stopped at the indicated time points; the cells were lysed, and TopoIIβK production was analyzed by Western blot using a mouse anti-TopoIIβK antibody. The infection in the absence of E64d was used as a negative control. The results show significant inhibition of TopoIIβK production by E64d, suggesting that TopoIIβK may be processed from gp120 by cysteine proteases. Graphical representation of densitometric analysis of TopoIIβK protein levels using ImageJ software. Data represent the mean ± SD from 3 independent experiments. A paired t-test was performed for group comparisons. Exact *P* values are indicated in the figure ( *P* values: *<0.05, **<0.01, ***<0.005, and ****<0.001).

We further evaluated the effect of E64d on TopoIIβK formation during HIV-1 infection. SupT1 cells were preincubated with 10 µM E64d for 1 h, then infected with HIV-1_NL4-3_, and infection was terminated at the indicated time points. The formation of TopoIIβK in infected cells was analyzed from 30 min p.i. up to 96 h p.i. The results showed that TopoIIβK levels significantly decreased in the presence of E64d from 30 min p.i. and were completely inhibited at later time points (Fig. 7B). These results establish that during infection, TopoIIβK may be produced from gp120 via cysteine protease activity within endolysosomes.

To investigate the role of cathepsin L in TopoIIβK generation, the cells were treated with the cathepsin L inhibitor Z-FY-CHO (10 µM; Supporting data Fig. S7) during HIV-1 envelope transfection. In the untreated control, a 72-kDa TopoIIβK band was readily detected and increased over time in transfected cells (Fig. 8A). In contrast, treatment with Z-FY-CHO markedly reduced TopoIIβK formation throughout the time course of transfection (Fig. 8A), thus indicating that inhibition of cathepsin L specifically suppresses TopoIIβK generation.

**Figure 8.**
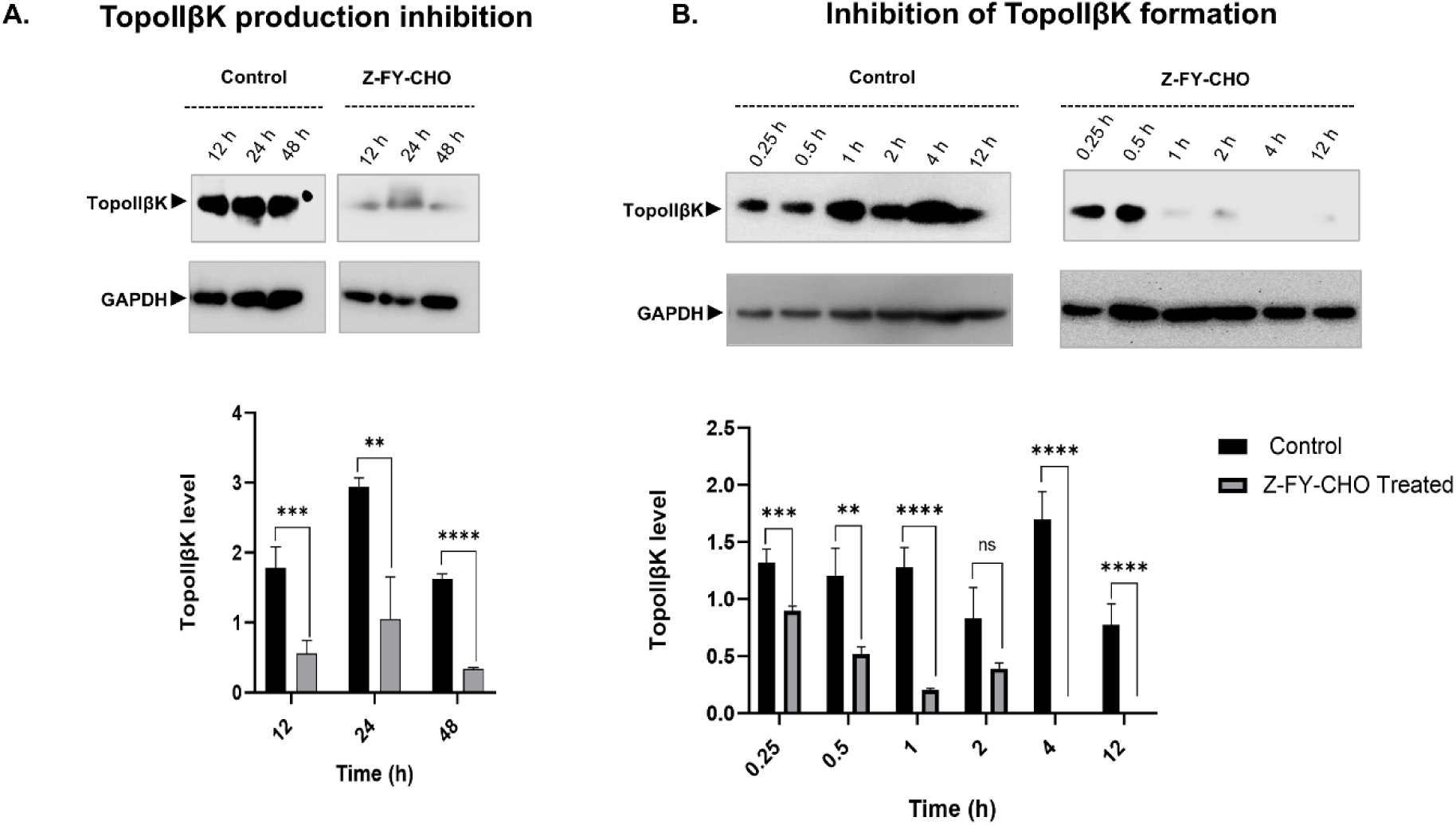
TopoIIβK may be processed from gp120 by cathepsin L. **(A)** TZM-bl cells were pre-incubated for 1 hour with the cathepsin L-specific inhibitor Z-FY-CHO, and then transfected with the subtype B gp120 plasmid (pTRO11). The cells were harvested at the indicated time points, and the production of TopoIIβK was analyzed by Western blot using a mouse anti-TopoIIβK antibody. Transfection in the absence of Z-FY-CHO is included as a negative control. The results show a highly significant reduction of TopoIIβK production in Z-FY-CHO-treated cells. **(B)** TZM-bl cells were pre-incubated with 10 µM Z-FY-CHO for 1 hour. These cells were infected with HIV-1 NL4-3 (2 MOI). Infection was stopped at the indicated time points, the cells were lysed, and TopoIIβK production was analyzed by Western blot using a mouse anti-TopoIIβK antibody. The infection in the absence of Z-FY-CHO was used as a negative control. The results show a complete inhibition of production of TopoIIβK by Z-FY-CHO, suggesting TopoIIβK may be processed from gp120 by cathepsin L. Graphical representation of densitometric analysis of TopoIIβK protein levels using ImageJ software. Data represent the mean ± SD from 3 independent experiments. A paired t-test was performed for group comparisons. Exact *P* values are indicated in the figure (*P* values: *<0.05, **<0.01, ***<0.005, and ****<0.001).

A similar reduction in TopoIIβK production was observed during HIV-1 infection in the presence of Z-FY-CHO (10 µM). Densitometric analysis in Fig. 8B confirmed a pronounced decrease in TopoIIβK levels in inhibitor-treated infection compared with untreated controls, with the strongest inhibition occurring during the early stages of infection (15–30 min) and persisting inhibition at later time points. These findings demonstrate that cathepsin L activity is required for efficient generation of the TopoIIβK. The results of experiments using E64d and Z-FY-CHO clearly demonstrate that TopoIIβK is produced by proteolytic cleavage of the substrate gp120 by cysteine proteases, most likely cathepsin L.

## Discussion

Viruses exploit cellular replication machinery; EBV, HBV, HSV, Ebola virus, and KSHV, alongside HIV, exploit cellular proteins [15–20]. HIV employs a rich arsenal of viral auxiliary proteins, in combination with cellular factors, in an exquisitely coordinated manner to complete the replication cycle and propagate [21,22]. Among the TopoII isoforms, TopoIIβ plays multiple roles during various stages of HIV-1 replication [8]. TopoIIβ activity is enhanced by phosphorylation during HIV-1 infection [10,11,23]. TopoIIβK inhibition impairs reverse transcription, suggesting that TopoIIβ phosphorylation is important for the strand transfer reaction [14]. The MALDI/ToF analysis of TopoIIβK identified peptides with envelope-like similarity (Fig. 3). Further immunological analysis of TopoIIβK for the presence of gp120 epitopes showed its immunoreactivity to antibodies recognizing an amino-terminal spanning V1 and V2 loops of gp120 (Fig. 4). A recombinant clone derived from the N-terminal region of gp120 (precTopoIIβK) showed phosphorylation of 180 kDa TopoIIβ and was also sensitive to an antagonist of TopoIIβK. Taken together, these results confirm that TopoIIβK is derived from gp120.

The most interesting aspect is the immunoreactivity to 2909, an antibody recognizing the trimeric V2-V3 loop conformation in the gp120 trimer [24]. This indicates that an anti-TopoIIβK antibody raised against native TopoIIβK may be recognizing this epitope, which may not be accessible in monomeric gp120. Such a conserved structure may be common across subtypes [25, 26, 35]. Thus, TopoIIβK purified from the virus particle, as well as gp120 clones, are conserved among all subtypes of viruses. As reported in the literature [36], during post-infection, lysosomes appear to encapsulate gp120 and reposition it from the perinuclear membrane towards the cell membrane [37]. Indeed, we observed that TopoIIβK is localized in endolysosomes at 15 min of p.i. suggesting that TopoIIβK could be produced in endolysosomes during HIV-1 infection. Studies by Miyauchi et al. (2009) [38] showed that Dynamin- and Clathrin-mediated endocytosis are prerequisites for viral fusion. Additionally, endocytosis inhibitors (EIPA, Pitstop2, Dynasore) have been shown to inhibit endocytic uptake and internal fusion of HIV-1, thereby reducing viral infection [39]. In the current study, we analyzed whether TopoIIβK is processed from gp120 by cysteine proteases. This analysis was conducted using the synthetic cysteine protease inhibitor E64d and the cathepsin L inhibitor, Z-FY-CHO. The production of TopoIIβK was monitored while cysteine protease activity was suppressed with E64d and Z-FY-CHO. The results in Fig. 7 showed that infection in E64d-treated Sup T1 cells significantly affected TopoIIβK production. Also, envelope transfection in cells treated with E64d affected TopoIIβK production. Furthermore, inhibition of cathepsin L with a specific inhibitor confirmed that TopoIIβK production was abrogated both in gp120-transiently expressing cells and during infection (Fig. 8). These observations unequivocally demonstrate the role of endolysosomal cysteine proteases, specifically cathepsin L, in processing the envelope to form TopoIIβK (Fig. 9). In studies analyzing the role of endosomal internalization in HIV-1 infection using gene knockdown, it was shown that the virus may utilize multiple pathways, including endocytosis [4]. These studies confirm our current observation that endoproteases are involved in envelope processing. Further, in the lysosomal environment, gp120 is exposed to several enzymes; among these, cysteine proteases such as cathepsin L and B are prominent [30,31,32]. Several reports have indeed shown the presence of a ∼70 kDa protein in viral particles, infected cells, and envelope-expressing cells [32,40–42], as well as in proteolytic analyses of gp120/envelope [41,43–47]. But we are not sure whether the 70 kDa reported in these publications corresponds to TopoIIβK. Also, further studies are required to elucidate the role of the ∼50-kDa residual envelope polypeptide reported in the literature [40–42]. Based on these observations, we propose that during HIV-1 entry, the gp120 encapsulated in the endolysosome may be processed to form TopoIIβK in infected cells, thereby participating in various stages of viral replication (Fig. 9).

**Fig. 9.**
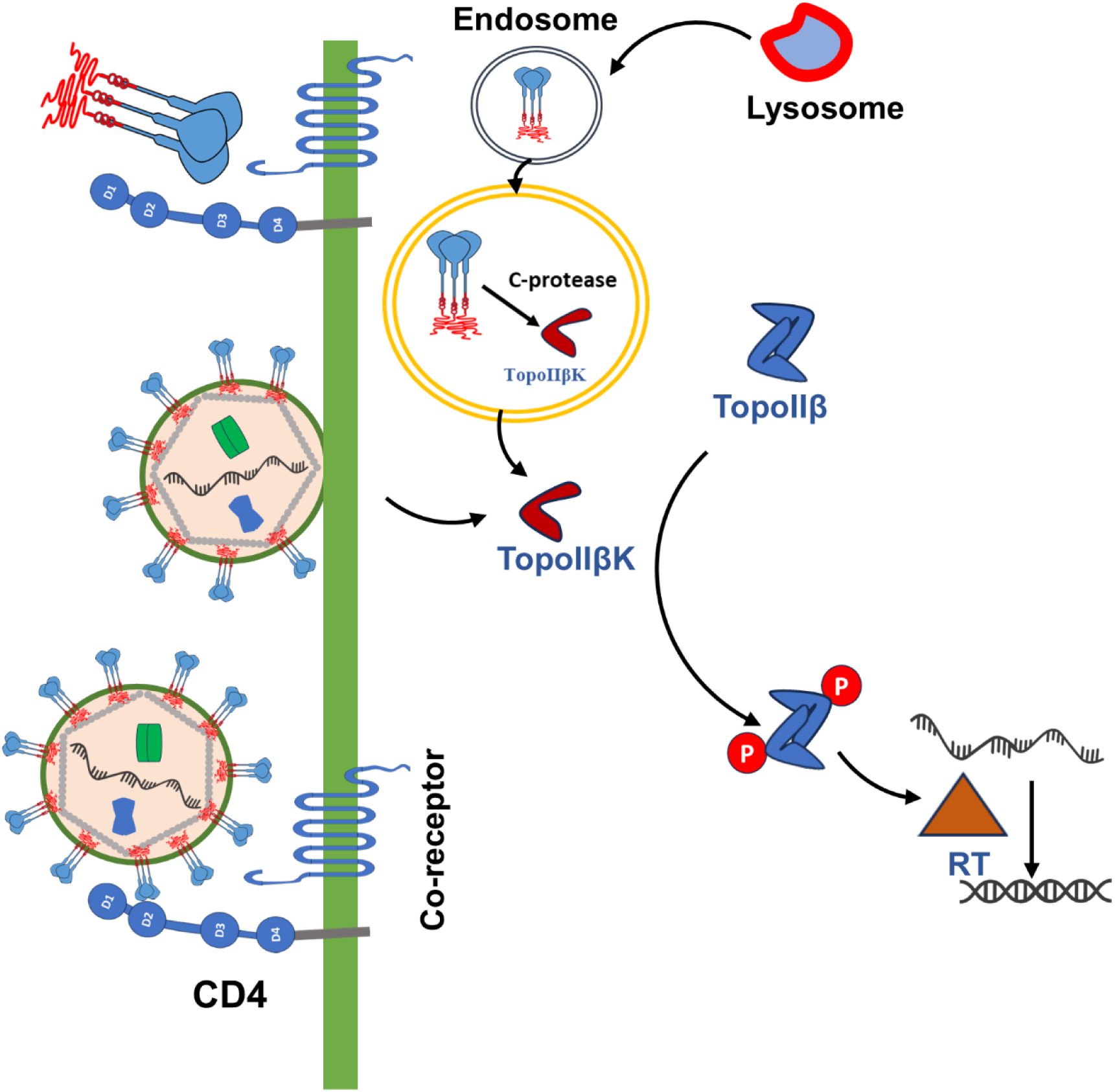
Graphical Abstract.

## Acknowledgments

Acknowledgments: HV (PMRF ID: 3702544) and RD (PMRF ID: 3702549) are PMRF Fellows. SM is supported by a DBT Senior Research Fellowship (No. DBT/2017/UOH/90). The authors thank Ms. Monica Kannan (RTGR-UoH Proteomics Facility) for MALDI-TOF experiments and analysis. The infrastructure support was provided by the IoE, DST–FIST, and DBT Builder programs (BT/INF/22/SP41176/2020). We thank Prof. Udaykumar Ranga, JNCSR, and Prof. Sharmistha Banerjee for providing pVSV-G.

Reagents were obtained through the NIH HIV Reagent Program, Division of AIDS, National Institute of Allergy and Infectious Diseases, National Institutes of Health: Env clones p398F1 (A), pTRO11 (B; Cecilia Graziosi), pCE1176 (C; Center for HIV/AIDS Vaccine Immunology (CHAVI)), and p246F3 (AC recombinant; Michael Hoelscher and Leonard Maboko), pBJOX2000 (CRF_BC; Hao Wu and Ning Li), HIV subtype panels (Dr. Victoria Polonis and Dr. Nelson Michael), and HIV-1 gp120 monoclonal antibodies (ID6 from Dr. Kenneth Ugen and Dr. David Weiner; A32-AAA and CH58 from the Duke Human Vaccine Institute) and cell lines SupT1 (Dr. James Hoxie); CHO_JR-FL_ (Dr. James Arthos); TZM-bl cell (Dr. John C. Kappes, Dr. Xiaoyun Wu, and Tranzyme Inc.).

## Author Contributions

A.K. Kondapi designed experiments and analyzed data.

H. Verma: Kinetics of TopoIIβ production in envelope clones, transfected cells, and different subtype-infected cells. TopoIIβK is characterized by a panel of anti-gp120 antibodies. Performed TopoIIβK localization in the endolysosome. Performed experiments using protease inhibitors in gp120-transfected cells and HIV-1 infected cells.

S. Mukhopadhyay: TopoIIβK is produced in HIV-1_NL4-3_ gp120- and gp160-transformed cells. Showed TopoIIβK is recognized by anti-gp120 antibody ID6. Performed experiments using a pseudo-virus.

L.S. Balakrishna: Showed TopoIIβK is produced in envelope-expressing CHO_JR-FL_ cells and performed MALDI TOF/TOF studies.

R. Das: Prepared anti-TopoIIβK antibody in a mouse model.

A. Kumar: Performed phosphorylation studies.

Sangeetha Balasubramanian: Sequence analysis of MALDI detected peptide.

H. Verma, S. Mukhopadhyay, and A.K. Kondapi: Compiled and formatted results & prepared manuscript.

## Data availability

All raw data supporting the findings of this study are available within the paper and its Supplementary Information.

## Abbreviations

HIV: Human Immunodeficiency Virus
E64d: (2*S*,3*S*)-3-[[[(1*S*)-3-Methyl-1-[[(3-methylbutyl)amino]carbonyl]butyl]amino]carbonyl]- 2-oxiranecarboxylic acid ethyl ester
Rab: Ras-associated binding
gp120: Glycoprotein 120
Tat: Trans-activator of Transcription
TopoIIβK: Topoisomerase II beta Kinase
MAP: Mitogen-activated protein
PBMCs: Peripheral blood mononuclear cells
MOI: Multiplicity of Infection
MALDI/TOF: Matrix-Assisted Laser Desorption Ionization/Time of Flight
HRP: Horseradish Peroxidase
NCBI: National Center for Biotechnology Information
MHC: Major Histocompatibility Complex
CHO_JR-FL_: Chinese Hamster Ovary cells expressing the envelope protein of the JR-FL strain of HIV-1
CRF: Circulating Recombinant Form of HIV-1
HEK: Human Embryonic Kidney
EBV: Epstein-Barr Virus
HBV: Hepatitis B Virus
HSV: Herpes Simplex Virus
KSHV: Kaposi Sarcoma-Associated Herpesvirus
EIPA: 5-(N-ethyl-N-isopropyl)-Amiloride
Z-FY-CHO: Z-Phe-Tyr-Aldehyde

