## Supplementary Data for "Topoisomerase IIβ Kinase: A protein processed from HIV-1 envelope by cysteine proteases"

##### **This file includes:**

Tables S1 to S5
Supporting text
Figures S1 to S8

**Table S1.** Details of HIV-1 viral isolates used in the study.

| **Subtype/CRF** | **Isolate** | **Country of origin** | **Contributor** |
| --- | --- | --- | --- |
| A | 99KE_KNH108 8 | Kenya (KE) | Dr. Victoria Polonis |
| B | 90US_873 | United States of America (USA) | Dr. Victoria Polonis |
| C | 94IN_20635-4 | India (IN) | Dr. Smita Kulkarni |
| C | 93IN101 | India (IN) | Dr. Robert Bollinger |
| D | 98UG_57128 | Uganda (UG) | Dr. Victoria Polonis |
| AE | 96TH_M02138 | Thailand (TH) | Dr. Victoria Polonis |

**Table S2.** Details of HIV-1 infectious clone and Envelope clones used in this study.

| **Subtype** | **ARP Reagent name** | **Genebank accession ID/Catalog No.** | **Country of origin** | **Contributor** |
| --- | --- | --- | --- | --- |
| A | p398F1 env | HM215312 | Tanzania | Michael Hoelscher and Leonard Maboko |
| B | HIV-1 pNL4-3 | AF324493.2 | - | Dr. Nathaniel Landau and Aaron Diamond. |
| B | pTRO11 env | AY835445 | Italy | Cecilia Graziosi |
| C | pCE1176 env | FJ444437 | Malawi | Center for HIV/AIDS Vaccine Immunology (CHAVI) |
| AC recombinant | p246F3 env | HM215364 | China | Michael Hoelscher and Leonard Maboko |
| CRF07_BC | pBJOX2000<br>env | HM21 5364 | China | Hao Wu and Ning<br>Li |
| pNL4-3ΔEnv | - | 3418 | - | Dr. Nathaniel<br>Landau |
| pVSV-G | - | 8454 | - | Bob Weinberg |

**Table S3.** Details of anti-GP120 monoclonal antibodies used in this study.

| **Antibody** | **ARP No** | **Epitope** | **Contributor** |
| --- | --- | --- | --- |
| ID6 | 2343 | Epitope corresponds to 86-100 Amino Acids | K Ugen, D Weiner |
| 2909 | 12141 | V2-V3 quaternary neutralizing epitope | S Zolla-Pazner |
| CH58 | 12550 | V2 epitope | B Haynes, H Liao, Duke Human Vaccine Institute, Duke University Medical Center |
| PG9 | 12149 | Quaternary structure of V1V2. | International AIDS Vaccine Initiative |
| 35022 | 12586 | Binds to the interface between gp120 and gp41 | J Huang, M Connors |
| 3BNC117 | 12474 | CD4 binding site of Envelope | M Nussenzweig |
| A32-AAA | 12554 | Recognizes the C1 region of HIV-1 envelope | B Haynes, H Liao, Duke Human Vaccine Institute, Duke University Medical Center |
| F425 | 7624 | CD4i epitope of gp120 | M Posner, L Cavacini |

**Table S4.** Details of Cell lines obtained from NIH

| **Cell Line** | **Isolate Name/ Origin** | **Contributor** |
| --- | --- | --- |
| SupT1 | Non-Hodgkin's T-cell Lymphoma | Dr. James Hoxie |
| CHOJR-FL | Chinese Hamster Ovary cells expressing the envelope protein of the JR-FL strain of HIV-1 | Dr. James Arthos |
| TZM-bl | Derived from the JC53 Cell by introducing integrated copies of the luciferase and β-galactosidase genes | Dr. John C. Kappes, Dr. Xiaoyun Wu, and Tranzyme Inc. |
| HEK293T | Human Embryonic Kidney Cells | Dr. Robert B. DuBridge, Dr. Peter Tang, Dr. Han-Chao Hsia, and colleagues |

### Supporting information:

#### Proteomic analysis of TopoIIβK

A

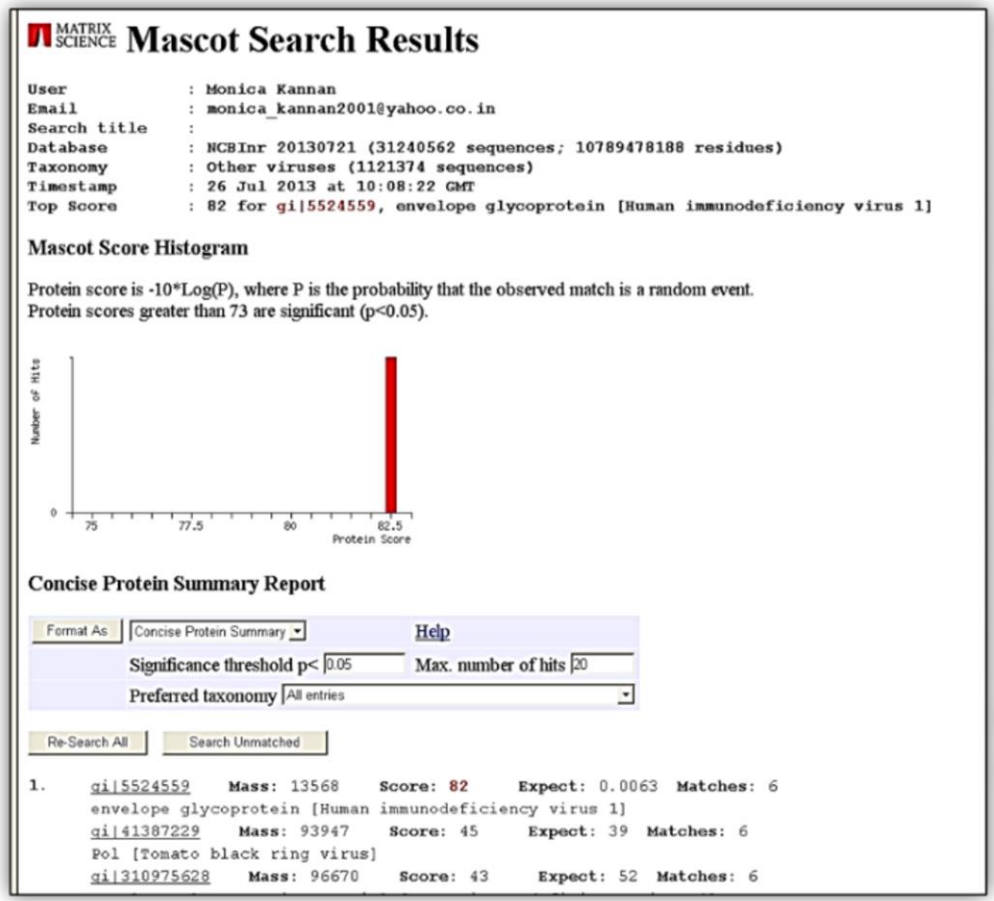

B

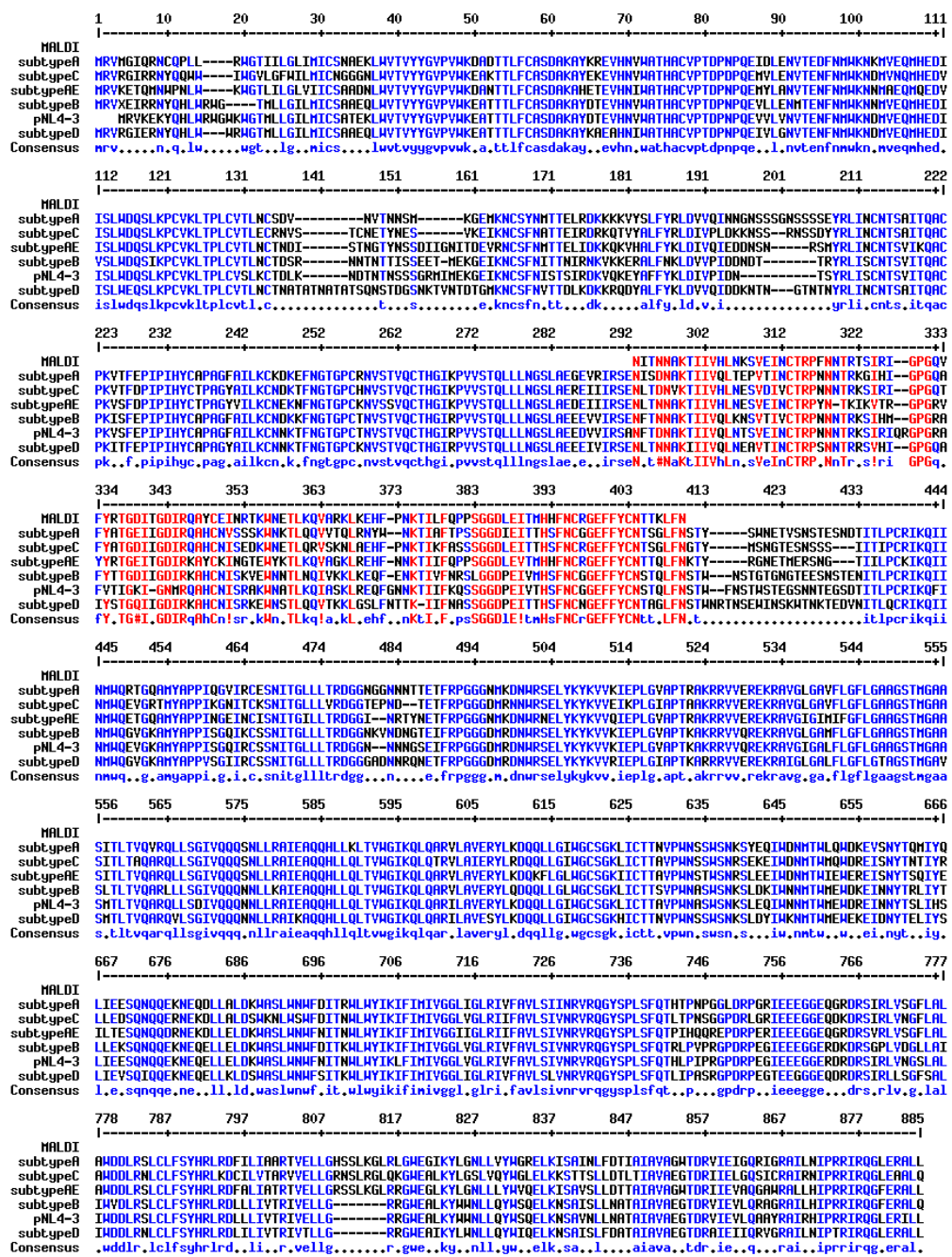

**Fig S1: Proteomic analysis of TopoIIβK. (A)** Mascot analysis of 72 kDa TopoIIβK protein purified from HIV-1 93In101 identified a single peptide homologous to the envelope glycoprotein of HIV-1. According to the mentioned envelope, a protein score greater than 73 is significant ($p < 0.05$), with a high likelihood score of 82. (B)

Multiple sequence alignment (performed with MultiAlin) results showed peptide homology with the envelope glycoprotein after trypsin digestion.

#### Host kinase cross-reactivity with TopoIIβK antibody

To check the role of the host kinase interacting with TopoIIβ during early time points of HIV-1 infection, reported TopoIIβK was evaluated for cross-reactivity with the reported cellular kinase CK1δ antibody. Host cell TZM-bl was infected with HIV-1 NL4-3(MOI 2), and cellular proteins were resolved on 8% SDS-PAGE along with the uninfected cell pellet and purified TopoIIβK and CK1δ.

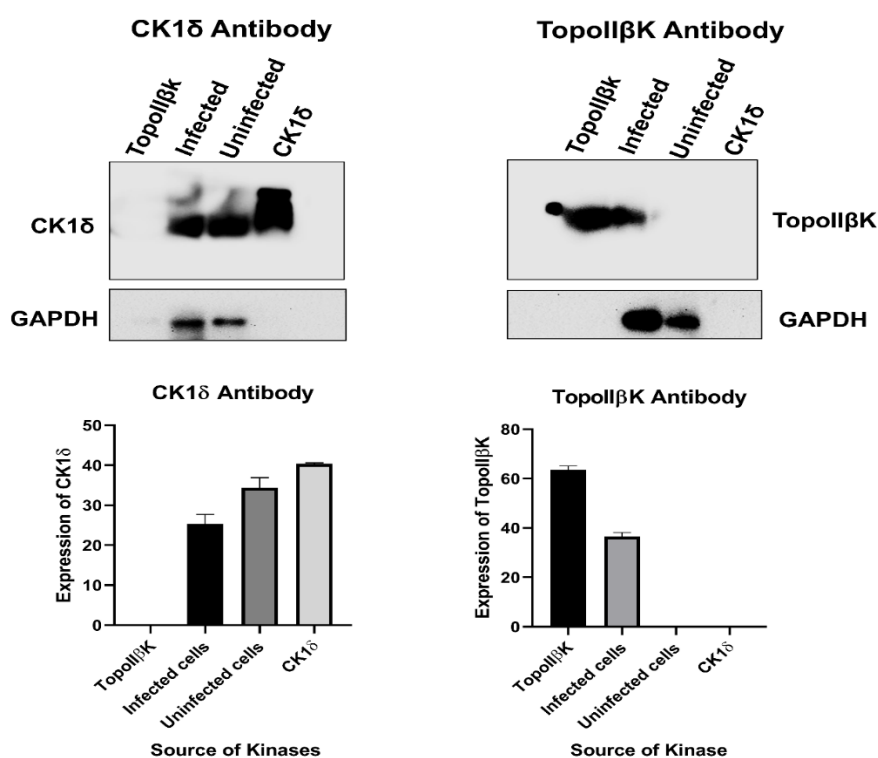

**Fig. S2: Host kinase cross-reactivity with TopoIIβK antibody.** HIV-1-infected TZM-bl cells were analyzed by Western blotting along with an uninfected cell pellet and purified TopoIIβK and CK1δ proteins. Results (**left panel**) depicted the presence of CK1δ in both uninfected cells and infected cells, while (right panel) TopoIIβK was

observed only in the infected cell pellet. (**Bottom Panels**) depicted densitometric analysis of the respective blots. Data represent the mean ± SD from three independent experiments performed in triplicate ($n = 3$)

### TopoIIβK purification from HIV-1 subtypes

TopoIIβK was purified from HIV-1 subtype 99KE\_KNH1088 (Subtype A), 90US\_873 (Subtype B), 94IN\_20635-4 (Subtype C), and 98UG\_57128 (Subtype D) by lysing the virus, followed by Mono Q anion exchange chromatography (as described in Methodology). The purified TopoIIβK was visualized by resolving it on a 10% SDS-PAGE gel and staining with Coomassie.

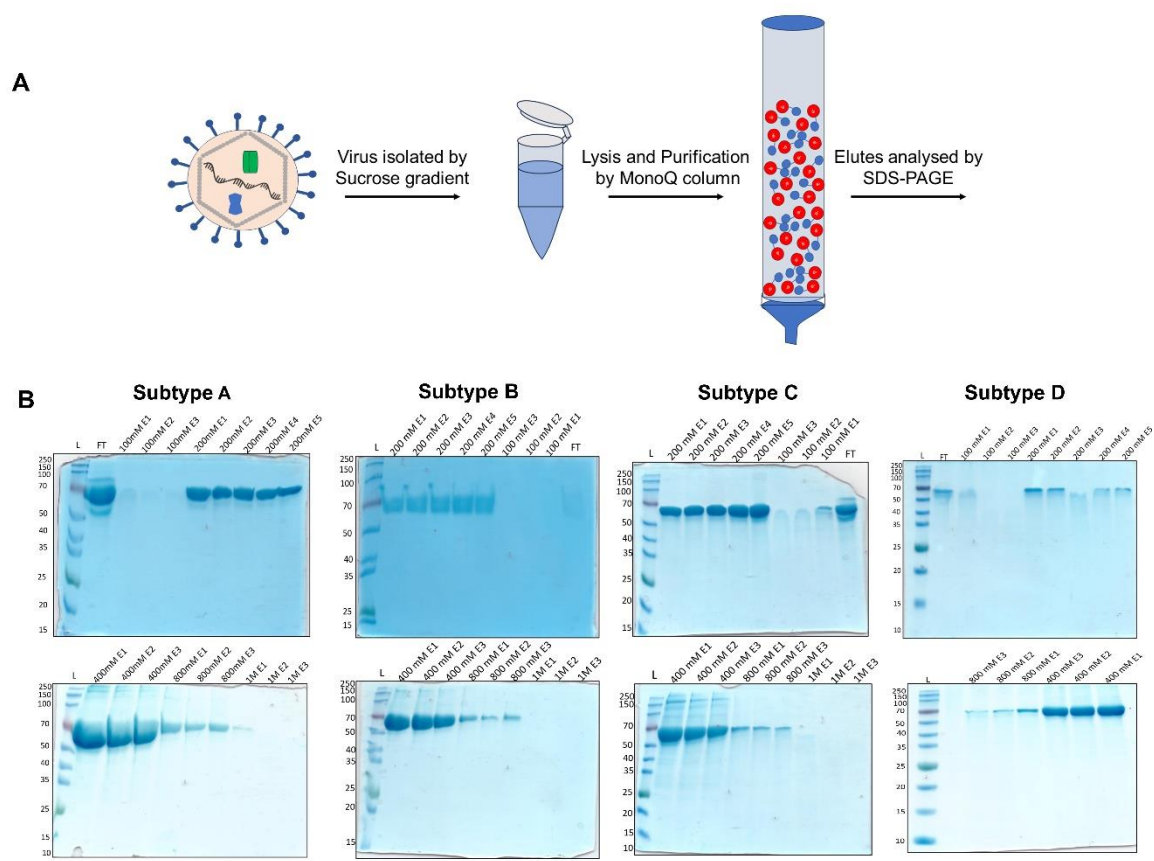

**Fig S3. (A)** Graphical representation of TopoIIβK purification from HIV particles. **(B)** SDS-PAGE analysis of HIV TopoIIβK purified by Mono-Q ion exchange chromatography. Subtype A shows the highest production of 72 kDa in the elution fraction at 200 mM and 400 mM. Subtype A shows maximum production of 72 kDa in the elution fraction at 400 mM. Subtype C shows the highest production, at 72 kDa, in the elution fractions at 200 mM and 400 mM. Subtype D shows a maximum production of 72 kDa at 400 mM in the elution fraction. Data represent the mean ± SD from three independent experiments performed in triplicate ($n = 3$).

##### gp120 clone expression HEK293T cells:

HEK293T cells at 70% confluency (1.8 million cells in a 60 mm cell culture dish) were transfected with Lipofectamine 3000 according to the manufacturer's guidelines using HIV-1 envelope clones (Table S2). Cells were fixed with 4% formaldehyde 24 hours post-transfection, followed by permeabilization and probed with mouse anti-TopoIIβK antibody and anti-mouse secondary antibody tagged with Alexa Fluor 594 (#A-11062).

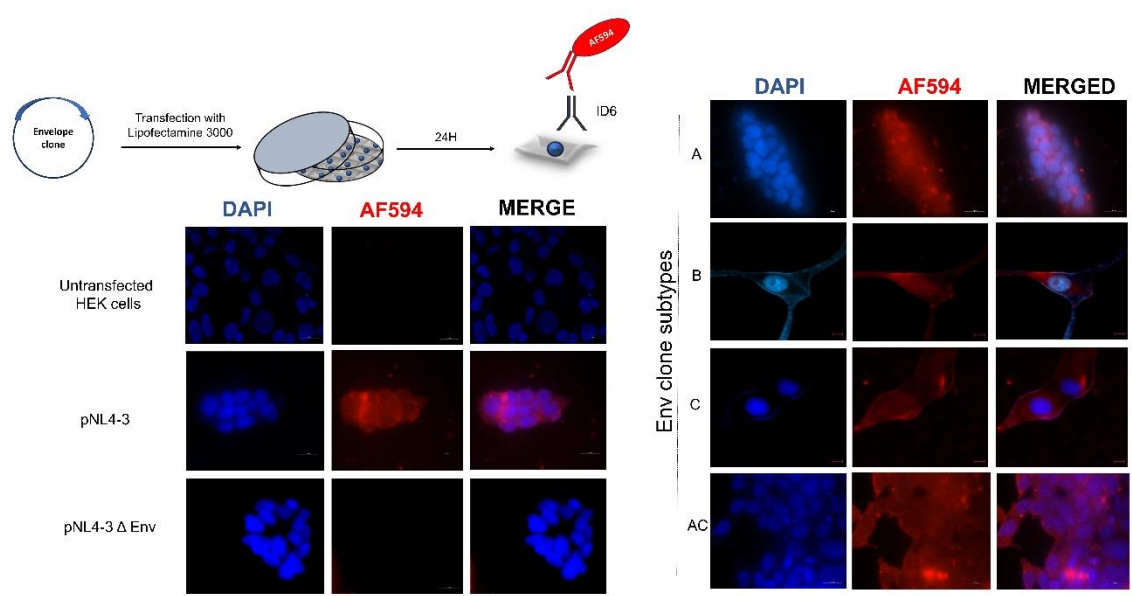

**Fig. S4: Envelope clone expression.** (A) Graphical representation of transfection-mediated expression. (B) Immunofluorescence studies of a gp120 clone expressing HEK293T cells showing recognition by anti-gp120 antibody ID6 and anti-mouse secondary antibody tagged with AF594, confirming transfection-mediated expression. Images were captured at 63x magnification using a Carl Zeiss Fluorescence Microscope and AxioCam 3.0.

#### **Detection of kinase activity by ADP-Glo assay**

Kinase activity was quantified using the ADP-Glo Kinase Assay kit (Promega), a luminescence-based assay that measures ADP generated by kinase-catalyzed phosphotransfer as a surrogate for enzymatic activity. Briefly, during the kinase reaction, ATP is consumed as the phosphate donor and converted to ADP upon phosphorylation of the substrate. The addition of the ADP-Glo reagent terminates the kinase reaction and depletes any residual unconsumed ATP. Subsequent addition of the kinase detection reagent converts the newly formed ADP back into ATP, which is then quantified using a coupled luciferase/luciferin reaction that generates a luminescent signal proportional to ATP (and hence ADP) concentration. Luminescence was recorded using a luminometer with an integration time of 0.5–1.0 s.

#### **Generation of an ADP standard curve**

An ADP standard curve was generated to convert luminescence readings into pmol of ADP produced. Ultrapure ATP and ADP stocks supplied with the kit were diluted in $1\times$ kinase buffer to prepare 1 mL of 1 mM ATP and 500 μL of 1 mM ADP working stocks. A master mix series (Row A) was then prepared by combining the ATP and

ADP stocks at defined ratios corresponding to 0–100% ADP conversion (Table S5), and 100 μL of each mix was added per well in Row A of a 96-well white opaque plate.

A three-step, 10-fold serial dilution of Row A was performed to generate Rows B–D: 90 μL of 1× kinase buffer was added to each well of Rows B, C, and D, and 10 μL was serially transferred from Row A → B → C → D, mixing thoroughly at each step.

ADP-Glo reagent (equal volume to the reaction/standard volume) was then added to each well, and the plate was incubated for 40 min at room temperature to deplete residual ATP. Kinase detection reagent was subsequently added, and the plate was incubated for a further 30 min at room temperature to convert ADP to ATP and generate the luminescent signal. Luminescence (RLU) was recorded using a luminometer with an integration time of 0.5–1.0 s, and a standard curve was constructed by plotting RLU against known ADP concentration (or % ADP).

**Table S5.** Composition of the ADP: ATP standard mixtures (Row A)

| Row |  | ADP % | ATP % |
| --- | --- | --- | --- |
| A1 |  | 100 | 0 |
| A2 |  | 80 | 20 |
| A3 |  | 60 | 40 |
| A4 |  | 40 | 60 |
| A5 |  | 20 | 80 |
| A6 |  | 10 | 90 |
| A7 |  | 5 | 95 |
| A8 |  | 4 | 96 |
| A9 |  | 3 | 97 |
| A10 |  | 2 | 98 |
| A11 |  | 1 | 99 |
| A12 |  | 0 | 100 |

### Measurement of kinase reaction products

Following the kinase reaction, an equal volume of ADP-Glo reagent was added to each sample, and the samples were incubated for 40 min at room temperature to terminate the reaction and deplete unconsumed ATP. Kinase detection reagent was then added, and samples were incubated for a further 30 min at room temperature to generate a luminescent signal proportional to the ADP produced. Luminescence was measured using a luminometer with an integration time of 0.5–1.0 s, and the amount of ADP generated in each reaction (pmol) was interpolated from the ADP standard curve.

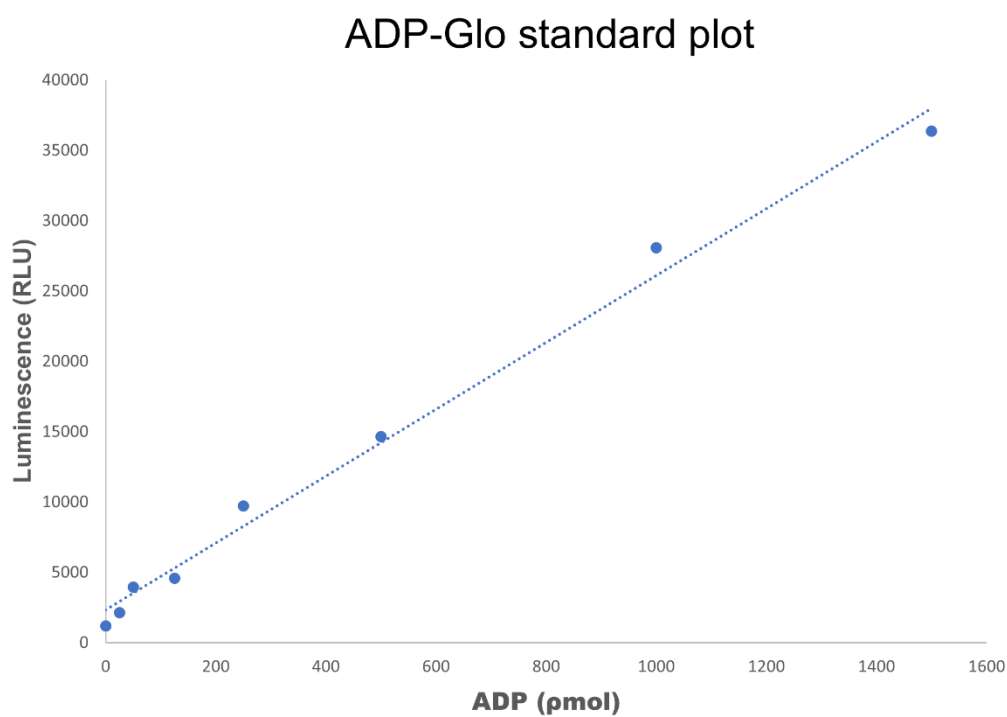

**Fig S5: Calibration curve for ADP-Glo assay.** Calibration curve generated using ATP/ADP standards supplied with the ADP-Glo® Kinase Assay kit. Luminescence (relative light units, RLU) was plotted against ADP concentration (pmol), and the resulting linear regression was used to interpolate ADP production in kinase reactions.

### Cytotoxicity Assay

Each well of a 96-well plate was seeded with $0.2 \times 10^6$ cells. Different concentrations of cysteine protease inhibitor (E64d) and Cathepsin L inhibitor (Z-FY-CHO) ($0.5 \mu\text{M}$, $1 \mu\text{M}$, $1.5 \mu\text{M}$, $2 \mu\text{M}$, $5 \mu\text{M}$, $10 \mu\text{M}$, and $20 \mu\text{M}$) were incubated for 24 hours. MTT solvent was incubated at 37 degrees Celsius at a concentration of $0.5 \text{ mg/ml}$ for 2 to 4 hours. $100 \mu\text{l}$ of DMSO was incubated at room temperature in the dark for 10-15 minutes. At $570 \text{ nm}$, the absorbance was determined using a microtiter plate reader.

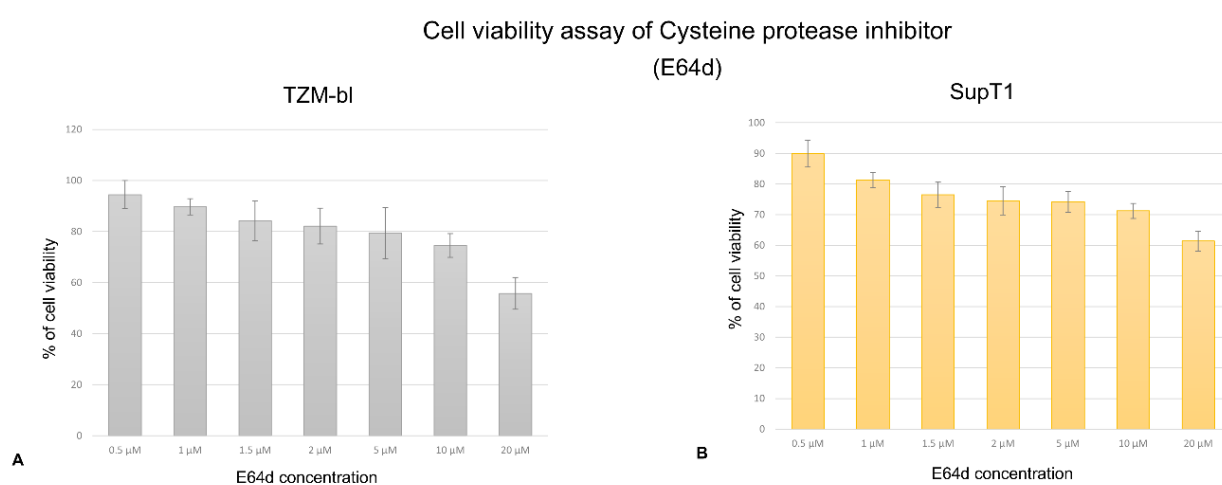

**Fig. S6: Cytotoxicity of E64d.** Cytotoxic effects of cysteine protease inhibitor E64d on cell lines **(A)** TZM-bl and **(B)** SupT1. Data represent the mean ± SD from three independent experiments performed in triplicate ($n = 3$).

Cell viability assay of cathepsin L inhibitor
(Z-FY-CHO)
TZM-bl

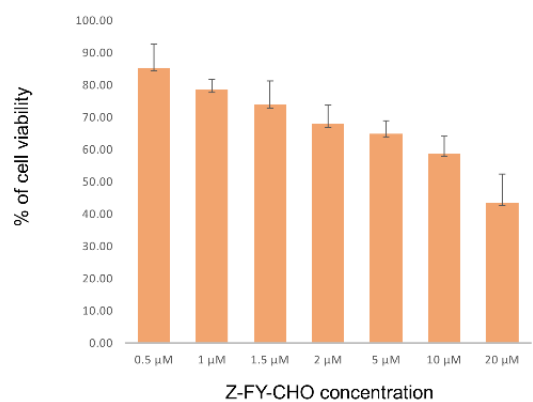

**Fig. S7: Cytotoxicity of Z-FY-CHO.** Cytotoxic effects of cathepsin L inhibitor (Z-FY-CHO) on cell line TZM-bl. Data represent the mean ± SD from three independent experiments performed in triplicate ($n = 3$).

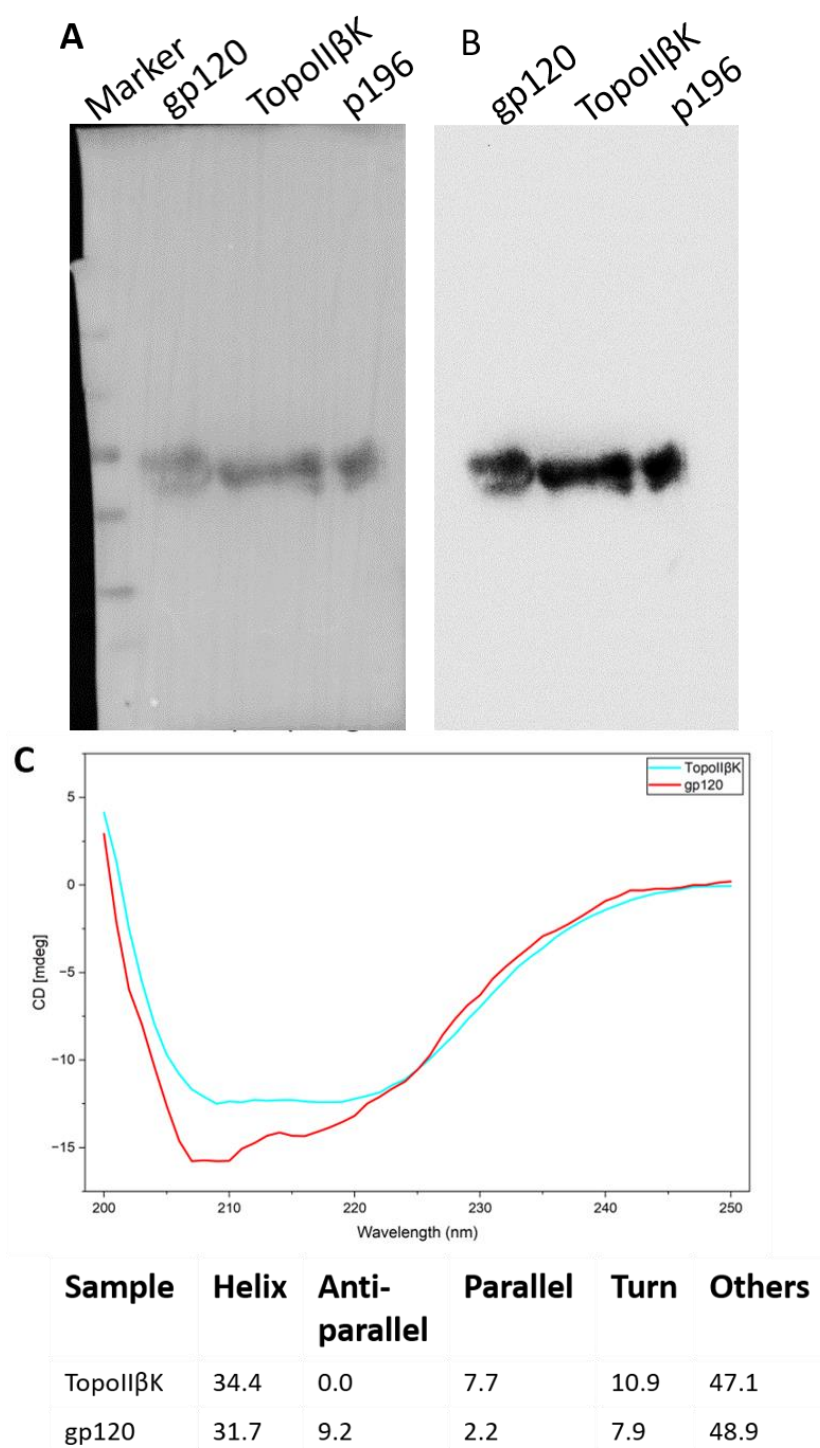

**Fig. S8. TopoIIβK-IgG does not recognize gp120.** The gp120 purified from ConA chromatography, TopoIIβK purified from MonoQ chromatography, and p196 (196 amino acids N-terminal polypeptide of gp120) were analyzed by Western blot using anti-TopoIIβK antibody (A: Blot with marker; B: Chemiluminescence image); the results show that the anti-TopoIIβK antibody recognizes only 72 kDa, not gp120.

Further, ConA-purified gp120 also co-elutes with TopoIIβK; B. CD spectrum of gp120 and TopoIIβK, which shows a significant change in conformation.
